# Loss of Ezh2 in the medial ganglionic eminence alters interneuron fate, cell morphology and gene expression profiles

**DOI:** 10.1101/2023.09.06.556544

**Authors:** Christopher T. Rhodes, Dhanya Asokumar, Mira Sohn, Shovan Naskar, Lielle Elisha, Parker Stevenson, Dongjin R. Lee, Yajun Zhang, Pedro P. Rocha, Ryan K. Dale, Soohyun Lee, Timothy J. Petros

## Abstract

Enhancer of zeste homolog 2 (Ezh2) is responsible for trimethylation of histone 3 at lysine 27 (H3K27me3), resulting in gene repression. Here, we explore the role of Ezh2 in forebrain GABAergic interneuron development. Loss of *Ezh2* increases somatostatin-expressing (SST+) and decreases parvalbumin-expressing (PV+) interneurons in multiple brain regions. We also observe fewer MGE-derived interneurons in the first postnatal week, indicating reduced interneuron production. Intrinsic electrophysiological properties in SST+ and PV+ interneurons are normal, but PV+ interneurons display increased axonal complexity in *Ezh2* mutant mice. Single cell multiome analysis revealed differential gene expression patterns in the embryonic MGE that are predictive of these cell fate changes. Lastly, CUT&Tag analysis revealed differential H3K27me3 levels at specific genomic loci, with some genes displaying a relative increase in H3K27me3 indicating they may be resistant to epigenetic modifications. Thus, loss of Ezh2 in the MGE alters interneuron fate, morphology, and gene expression and regulation.

## INTRODUCTION

Inhibitory GABAergic interneurons are a heterogeneous cell population with dozens of subtypes displaying distinct morphologies, connectivity, electrophysiology properties, neurochemical markers, and gene expression profiles^1–4^. Perturbation of interneuron development and inhibition is associated with a range of disorders including epilepsy, schizophrenia and autism^5–8^, and many disease-associated genes are enriched in prenatal immature interneurons and affect their development^9–11^. Forebrain interneurons originate from two transient structures in the embryonic ventral forebrain, the medial and caudal ganglionic eminences (MGE and CGE, respectively), and mature over the course of embryonic and postnatal development^1,3,4,12^. The MGE gives rise to distinct, non-overlapping interneuron subtypes, parvalbumin-(PV+) and somatostatin-expressing (SST+) interneurons (fast-spiking (FS) and non-fast spiking (NFS) interneurons, respectively).

Several factors regulate initial interneuron fate decisions within the MGE, including gradients of diffusible cues, spatial location of progenitors, temporal birthdates and the mode of neurogenesis^4,13–20^. The advent of single cell sequencing technologies over the last decade has generated a transcriptional and epigenetic ‘ground truth’ in the ganglionic eminences in mice^16,19–23^, and more recently, in primates and humans^24–31^. With this baseline in place, researchers can better characterize how genetic and epigenetic perturbations affect the fate and maturation of GABAergic interneurons.

Epigenetic mechanisms play critical roles in gene expression during neurogenesis, and modifications of the chromatin landscape regulate cell state changes during neurodevelopment^32–35^. Alterations in epigenetic regulation can be associated with numerous neurodevelopmental disorders^36–38^. Enhancer of Zeste Homolog 2 (Ezh2) is the primary methyltransferase component of the Polycomb Repressive Complex 2 (PRC2) that is critical for trimethylation of histone 3 at lysine 27 (H3K27me3) resulting in gene repression^39–41^. *Ezh2* is an evolutionary conserved gene that is aberrantly overexpressed in several forms of cancerous tumors^42,43^. *EZH2* variants can lead to Weaver Syndrome, a complex disease with variable degrees of intellectual disability^44,45^, and dysregulation of H3K27me3 may be the primary driver in ataxia-telangiectasia^46^. *Ezh2* expression is enriched in neural precursor cells where it represses target genes crucial to cell fate decisions and, in concert with other epigenetic marks, generates a transcriptional memory of specific gene expression patterns through cell divisions^39,47,48^. Loss of *Ezh2* can lead to ectopic exiting of the cell cycle and premature neuronal differentiation^49–53^, neuronal migration defects^54,55^, altered neuronal fate^50,56,57^ and changes in neuronal morphology and cognitive defects^58,59^. Thus, Ezh2 is an important player in epigenetic regulation of neuronal fate and maturation, but a role for Ezh2 in forebrain GABAergic interneurons has not been explored.

We generated conditional *Ezh2* knockout (KO) mice to remove *Ezh2* from the MGE and observed an increase in SST+ and decrease in PV+ interneurons across multiple brain regions. These fate changes were due to *Ezh2* loss in cycling neural progenitors, as removing *Ezh2* in postmitotic cells did not alter interneuron fate. While the intrinsic physiology of MGE-derived interneurons in *Ezh2* KO mice was normal, fast-spiking cells displayed an increase in axonal length and branching. Fewer cortical MGE-derived interneurons were observed during the first postnatal week, which likely indicates decreased neurogenesis compared to WT mice. Single cell transcriptome analysis revealed an increase in *SST*-expressing cells and a decrease in PV-fated cells during embryogenesis, consistent with the fate changes observed in the adult. Lastly, while a global downregulation of H3K27me3 was observed in the *Ezh2* KO MGE, we observed a relative increase in H3K27me3 at specific genomic loci in the KO, indicating that global loss of *Ezh2* had a differential effect at specific loci. In sum, we demonstrate that loss of *Ezh2* disrupts H3K27me3, alters gene expression and cell proliferation in the MGE, which in turn disrupts the normal balance of SST+ and PV+ interneurons in the forebrain.

## RESULTS

### Loss of *Ezh2* alters MGE-derived interneuron fate in the cortex and striatum

To characterize the function of *Ezh2* in MGE-derived interneurons, we crossed *Nkx2.1-Cre^C/C^*;*Ezh2^F/+^*males with *Ai9^F/F^*;*Ezh2^F/+^* females to generate *Nkx2.1-Cre^C/+^*;*Ezh2*;*Ai9^F/+^* wildtype, heterozygous and knockout mice (hereafter WT, Het and KO mice, respectively). In these mice, *Ezh2* perturbation is restricted to MGE-derived cells in the telencephalon, and these cells also express tdTomato (Tom+). We first confirmed that *Ezh2* expression is strongly downregulated in the MGE of KO mice (Fig. 1A). Since Ezh2 is critical for tri-methylation at histone H3K27, we also verified a significant reduction in H3K27me3 in the MGE in KO mice (Fig. 1B). To quantify this decrease, we performed Western Blots on MGE tissue from WT, Het and KO mice. We observed a 15% decrease of H3K27me3 signal in Hets and a 47% decrease in KO MGE compared to WT levels (Fig. 1C). Some of this remaining H3K27me3 in the KO MGE likely arises from the dorsal MGE where Cre expression is lacking in *Nkx2.1-Cre* mice^60^ (Fig. 1B), and from non-*Nkx2.1* lineage cells within the MGE (endothelial cells, LGE- and CGE-derived cells migrating through this region, etc.).

**Fig. 1.**
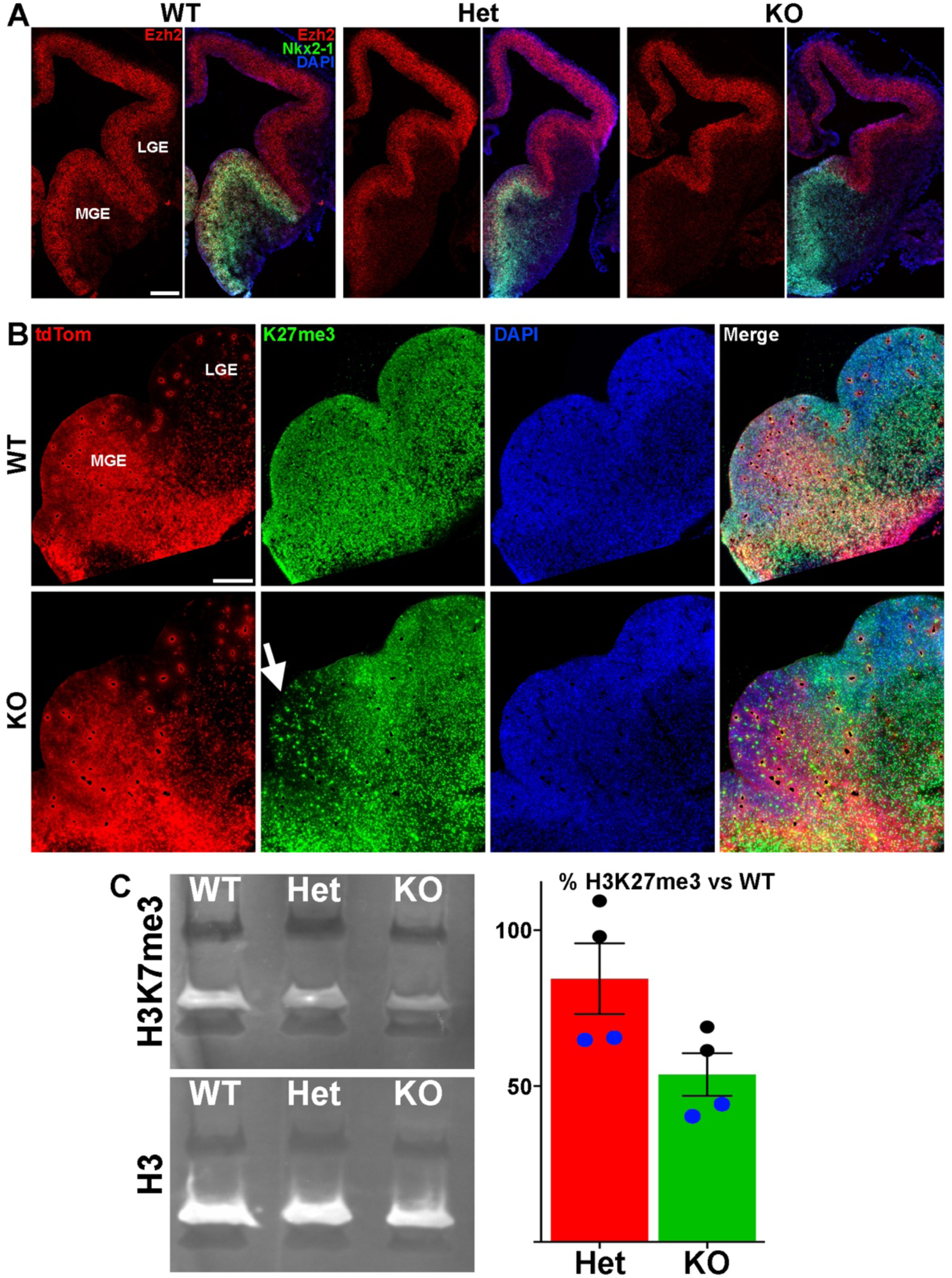
Loss of *Ezh2* and H3K27me3 in the MGE of *Ezh2* KO mice. **A.** *In situ* hybridizations of *Ezh2* (red) and *Nkx2.1* (green) in E12.5 brains of *Nkx2.1-Cre*;*Ezh2*;*Ai9* WT, Het and KO mice. **B.** Immunostaining in the MGE reveals a strong decrease of H3K27me3 (green) in the MGE of KO mice. **C.** Representative Western Blot gel showing H3K27me3 and H3 levels in WT, Het and KO MGE (left) and graph summarizing average H3K27me3 decrease in Het and KO MGE (right), with black and blue dots representing 2 different biological reps, with 2 technical reps each. Scale bars in A and B = 200 μm.

To determine if loss of *Ezh2* affects interneuron fate, we counted the number of SST+ and PV+ interneurons in the somatosensory cortex, hippocampus and striatum from WT, Het and KO mice (Fig. 2A). We observed a moderate but significant decrease in the density of Tom+ MGE-derived cortical interneurons in the cortex of KO mice. More striking, there was a significant increase in the density of SST+ interneurons and a corresponding decrease in the density of PV+ interneurons in the cortex of KO mice (Fig. 2B). This shift in cell fate is also apparent when examining the proportion of Tom+ cells that express either SST or PV (Fig. 2C).

**Fig. 2.**
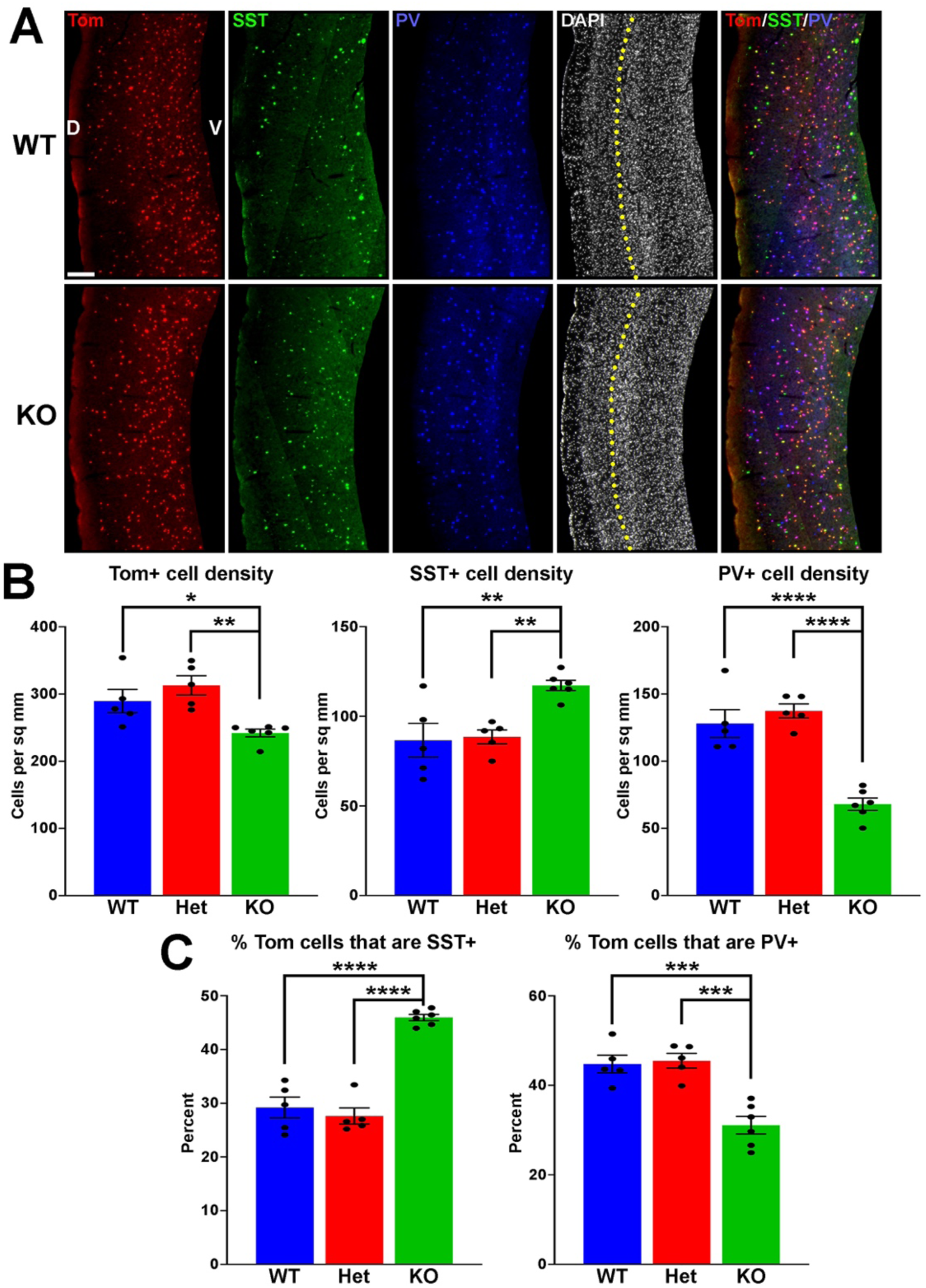
Changes in cortical interneuron fate in *Ezh2* KO mice. **A.** Representative images through the somatosensory cortex of P30 *Nkx2.1-Cre*;*Ezh2*;*Ai9* WT and KO mice stained for SST (green) and PV (blue). Scale bar = 100 μm. Yellow dotted lines indicate division between superficial (layers I-III) and deep (layers IV-VI) cortical layers defined by differential DAPI densities. D = dorsal, V = ventral. **B.** Graphs displaying the density of Tom+, SST+ and PV+ cells in WT, Het and KO mice. **C.** Graphs displaying the percent of Tom+ cells expressing SST or PV in WT, Het and KO mice. For all graphs, statistics are one-way ANOVA followed by Tukey’s multiple comparison tests: * = p ≤ .05, ** = p ≤ .005, *** = p ≤ .0005, **** = p ≤ .0001. n = 5 WT, 5 Het and 6 KO brains from a total of 4 different litters.

To examine if these differences were consistent throughout cortical layers, we divided the somatosensory cortex into superficial (II-III) and deep (IV-VI) layers based on DAPI staining. There was no significant decrease in the density of Tom+ cells between WT and KO mice in either the superficial or deep layers (Supplementary Fig. 1). The strongest effect in SST and PV fate changes was found in deep cortical layers, whereas superficial layers displayed a more moderate increase in SST+ and decrease in PV+ cells (Supplementary Fig. 1). Thus, loss of *Ezh2* in the MGE results in a slight reduction in total MGE-derived cortical neurons, with a significant increase in SST+ and decrease in PV+ cells, most notably in the deeper cortical layers.

MGE-derived SST+ and PV+ interneurons also populate the adult striatum. We observed a significant decrease in the density and proportion of PV+ interneurons in the striatum of KO mice, but no change in the total number of Tom+ or SST+ cells (Supplementary Fig. 2), indicating this decrease in the density of PV+ interneurons is observed in multiple brain regions.

### Alteration in both hippocampal interneurons and oligodendrocytes in *Ezh2* KO

The hippocampus contains a population of MGE-derived, neuronal nitric oxide synthase nNos-expressing (nNos+) neurogliaform and ivy cells that are not found in the cortex^61–63^. SST+, PV+ and nNos+ interneurons each make up ∼1/3 of MGE-derived hippocampal interneurons^64^. We characterized the densities and percentages of SST+, PV+ and nNos+ interneurons throughout the hippocampus (Fig. 3A-B). No significant differences were found in the densities of Tom+, SST+, PV+ or nNos+ interneurons in the whole hippocampus (Fig. 3C). However, there was a significant increase in the percentage of Tom+ cells that expressed SST or nNos in KO mice compared to WT (Fig. 3D).

**Fig. 3.**
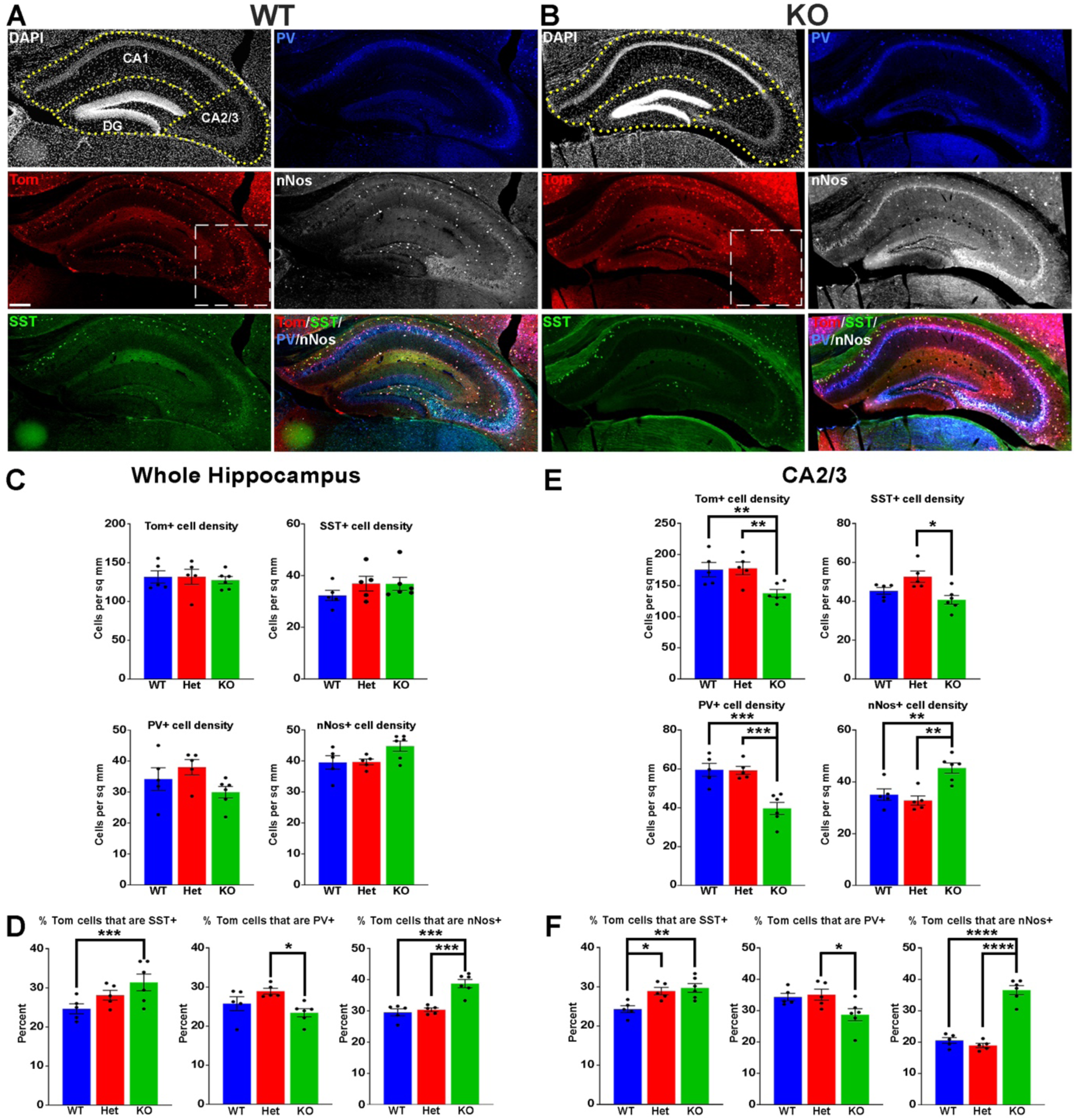
Changes in hippocampal interneuron fate in *Ezh2* KO mice. **A-B.** Representative hippocampus images of P30 *Nkx2.1-Cre*;*Ezh2*;*Ai9* WT (A) and KO (B) mice stained for SST (green), PV (blue) and nNos (white). Scale bar = 100 μm. Dotted white box indicates region blown up in Figure 4. **C-D.** Graphs displaying the density of Tom+, SST+, PV+ and nNos+ cells (C) and the percent of Tom+ cells expressing SST, PV or nNos (D) in the whole hippocampus of WT, Het and KO mice. **E-F.** Graphs displaying the density of Tom+, SST+, PV+ and nNos+ cells (E) and the percent of Tom+ cells expressing SST, PV or nNos (F) in the CA2/3 region of WT, Het and KO mice. All stats are one-way ANOVA followed by Tukey’s multiple comparison tests: * = p ≤ .05, ** = p ≤ .005, *** = p ≤ .0005, **** = p ≤ .0001. n = 5 WT, 5 Het and 6 KO brains, from 4 different litters.

Since the prevalence of interneurons differs between the CA1, CA2/3 and dentate gyrus (DG) regions of the hippocampus, we divided hippocampal sections into these three regions based on DAPI staining (Fig. 3A-B). There were almost no differences in densities or the proportion of subtypes in the CA1, only a slight but significant increase in the percentage of Tom+/nNos+ cells in the KO (Supplementary Fig. 3A). This increase in the percentage of nNos+ cells was also detected in the DG. More striking was a strong reduction of both PV+ cell densities and the percentage of Tom+/PV+ in the DG (Supplementary Fig. 3B).

The CA2/3 region displayed the greatest differences between WT and KO mice, many of which mimicked cell fate changes in the cortex. First, there was a decrease in the total density of CA2/3 Tom+ cells in the KO compared to WT (Fig. 3E). Second, there was a significant decrease in PV+ cell densities in the CA2/3 region of KO mice, and a corresponding increase in the percentage of Tom+/SST+ (Fig. 3C). Third, there was a significant increase in the density and percentage of nNos+ cells in CA2/3 region of KO mice compared to WT (Fig. 3C). Thus, loss of *Ezh2* in the MGE had both broad and region-specific effects on interneuron fate in the hippocampus: an increase in the proportion of MGE-derived nNos+ cells in all three hippocampal regions, an increase in the percentage of Tom+/SST+ cells specifically in the CA2/3 region (which resulted in a significant increase in the entire hippocampus), and a decrease in PV+ cell density in both CA1 and CA2/3. This decrease in PV cells was observed in the cortex, striatum and hippocampus.

While performing cell counts in the CA2/3 region, we observed Tom+ cell bodies that were too small to be interneurons, and we did not observe these cells in other brain regions (Fig. 4A). Counting these cells separately, we found a very strong reduction of these CA2/3-specific cells in the KO hippocampus (Fig. 4B). We stained WT hippocampal sections with various glia and microglia markers and found that many of these small Tom+ cell bodies were Olig2+, indicating that they are likely oligodendrocytes (Fig. 4C). This decrease in oligodendrocytes in *Ezh2* KO mice is consistent with findings demonstrating that loss of *Ezh2* can block or delay gliogenesis^65,66^.

**Fig. 4.**
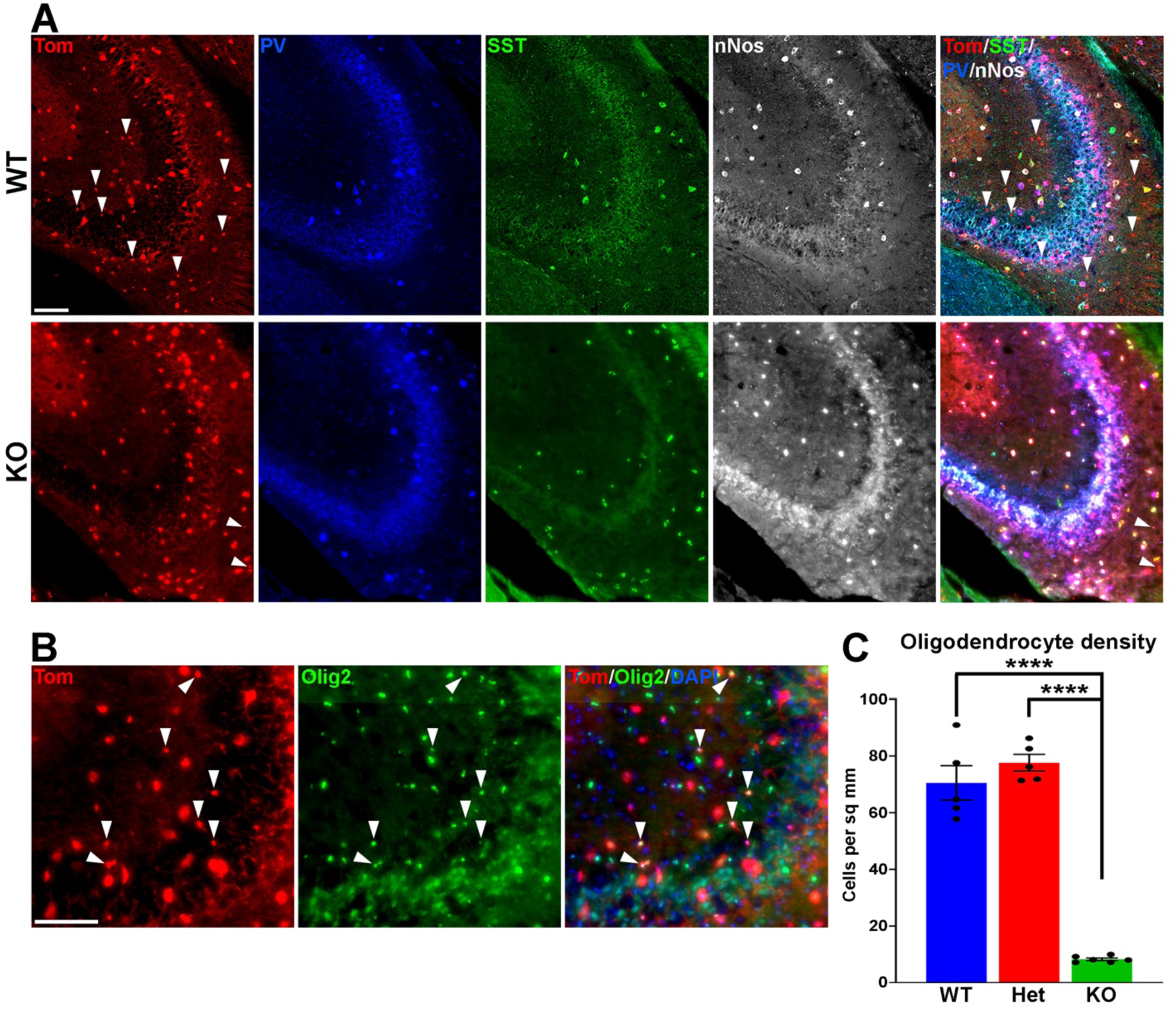
Loss of MGE-derived oligodendrocytes in CA2/3 in *Ezh2* KO mice. **A.** Representative images through the CA2/3 region of the hippocampus from P30 *Nkx2.1-Cre*;*Ezh2*;*Ai9* WT and KO mice stained for SST (green), PV (blue) and nNos (white). From boxed regions in Figure 3A-B. **B.** High magnification image of CA2/3 from *Nkx2.1-Cre*;*Ezh2*;*Ai9* WT mice showing many small Tom+ cells express the oligodendrocyte markers Olig2. Scale bars in A and B = 50 μm. **C.** Graph displaying the density of Tom+ oligodendrocytes in CA2/3 region of WT, Het and KO mice. All stats are one-way ANOVA followed by Tukey’s multiple comparison tests: **** = p ≤ .0001. n = 5 WT, 5 Het and 6 KO brains from a total of 4 different litters.

### Normal intrinsic properties but altered morphology of *Ezh2* KO interneurons

To characterize the intrinsic physiology of MGE-derived interneurons in KO mice, we performed patch clamp recording of layer V/VI Tom+ cortical cells in acute brain slices. Cells were classified as FS or NFS based on their intrinsic electrophysiological properties characterized under current clamp-recording. NFS cells had larger half-width, input resistance and membrane time constant/Tau, but smaller rheobase compared to FS cells. We analyzed action potential shapes, resting membrane potential, spike adaptation ratio, afterhyperpolarization (AHP) amplitude, input resistance and rheobase. There were no differences in intrinsic properties of FS or NFS cortical interneurons between WT and KO mice (Supplementary Fig. 4).

However, reconstructions of recorded cells did reveal morphological changes in FS cells. The axonal arbor of FS cells from KO mice were larger and more complex compared to WT cells (Fig. 5A). Sholl analysis revealed a significant increase in axon intersections, axon length and axon volume in FS KO cells, while there were no changes in dendritic arbors (Fig. 5B). This increased axonal arbor is similar to what was observed when trkB signaling was blocked in PV cells^67^. Thus, while intrinsic properties of Tom+ MGE-derived cortical interneurons were normal in *Ezh2* KO mice, FS cells displayed greater complexity in their axonal arbors compared to FS cells from WT mice.

**Fig. 5.**
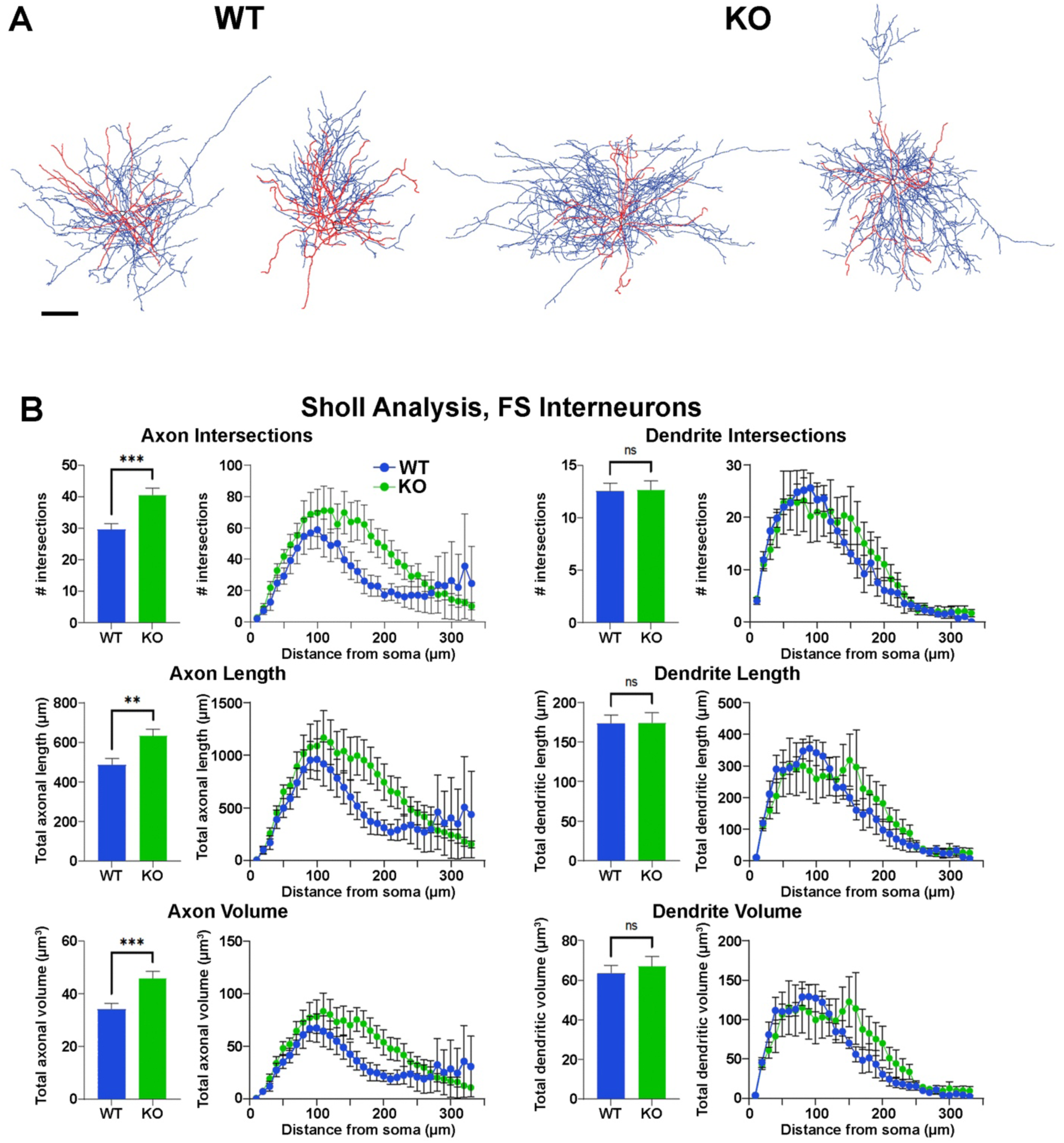
Increased axonal complexity of fast-spiking cortical interneurons in *Ezh2* KO mice. **A.** Representative morphological reconstructions of biocytin-filled FS cortical interneurons from *Nkx2.1-Cre*;*Ezh2*;*Ai9* WT and KO mice depicting axons (blue) and dendrites (red). Scale bar = 20 μm. **B.** Sholl analysis reveals increased axon intersections, axon length and axon volume in FS cortical interneurons from KO mice compared to WT littermates. No significant differences were found in the dendritic arbors. All statistics are two-way ANOVA followed by Holm-Sidak’s test: ** = p ≤ .005, *** = p ≤ .0005; n = 6 cells from 4 WT mice and 7 cells from 4 KO mice.

### Loss of *Ezh2* in cycling progenitors is required for cell fate changes

We next wanted to determine at what stage of development loss of Ezh2 results in cell fate changes. *Ezh2* is enriched in cycling progenitors throughout the embryonic brain (Fig. 1A), but Ezh2 may play a critical at other stages as well. To investigate this possibility, we generated *Dlx5/6-Cre*;*Ezh2^F/F^*;*Ai9* conditional KO mice in which loss of *Ezh2* is restricted to postmitotic neurons arising from the ganglionic eminences. We verified that *Ezh2* is still expressed in MGE ventricular zone cycling progenitors in *Dlx5/6-Cre* KO mice (Supplementary Fig. 5A). There were no differences in the densities or percent of SST+ and PV+ cells in the cortices of these *Dlx5/6-Cre* KO mice (Supplementary Fig. 5B), indicating that *Ezh2* is required in cycling MGE progenitors for proper interneuron fate and maturation.

A wave of programmed apoptosis occurs between the first and second postnatal weeks that eliminates ∼20-40% of cortical interneurons^68–71^. To determine if there were changes in the overall production of MGE-derived interneurons during embryogenesis, we counted the number of Tom+ cells in the cortex at P5, prior to programmed apoptosis. We found a significant decrease in the number of Tom+ cells in the KO cortex compared to WT at P5 (Fig. 6A-B). This finding supports the hypothesis that loss of *Ezh2* in cycling MGE progenitors decreases the overall production of MGE-derived cortical interneurons. The stronger decrease of Tom+ cells in the superficial layers is consistent with preemptive depletion of the progenitor pool, which would (1) primarily affect the later-born cells in the superficial layers, and (2) lead to more prominent loss of PV+ interneurons due to their bias production at later embryonic timepoints compared to SST+ cells^15^.

**Fig. 6.**
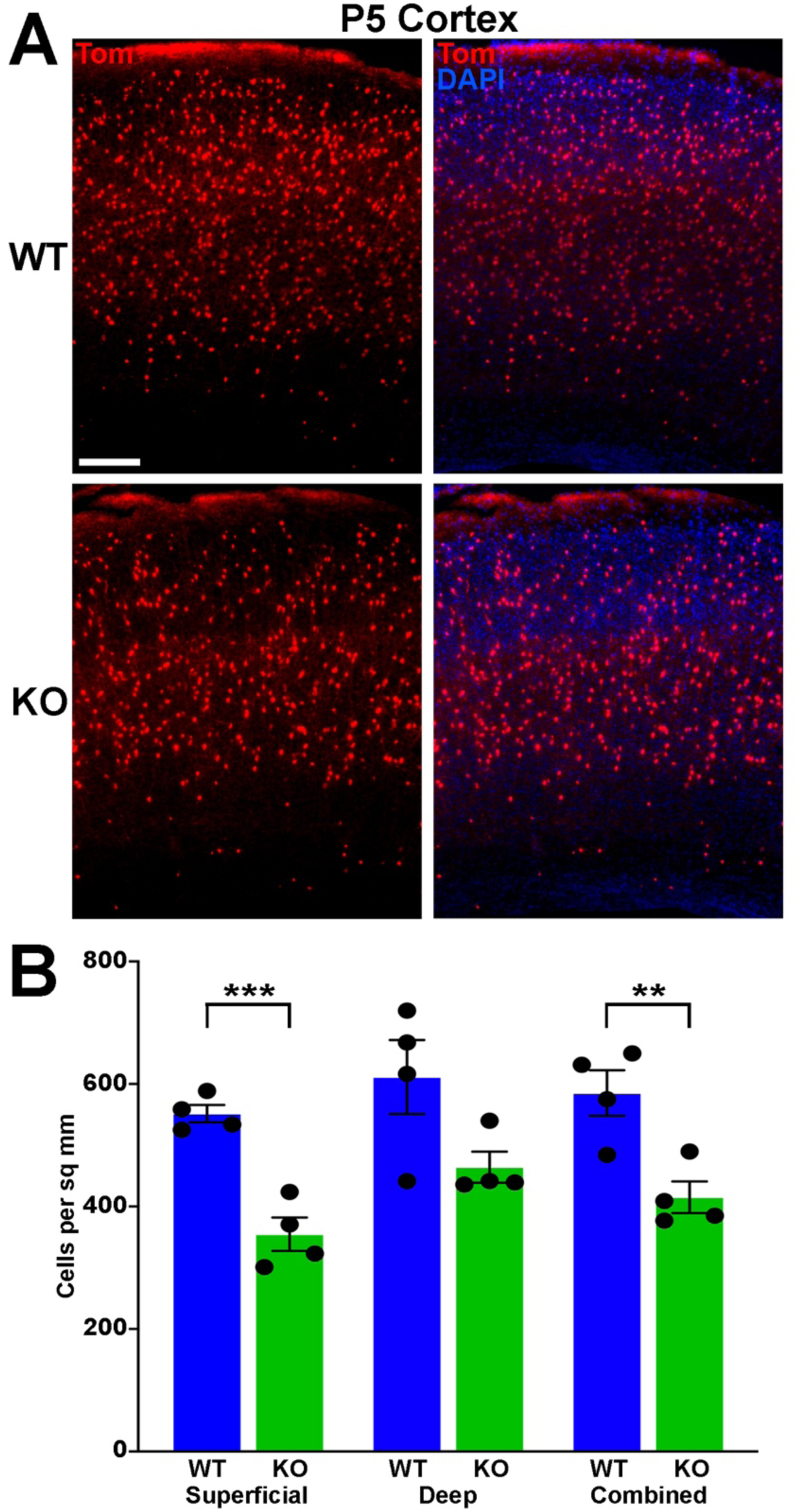
Significantly fewer MGE-derived cortical interneurons at P5 in *Ezh2* KO mice. **A.** Representative cortical images from P5 *Nkx2.1-Cre*;*Ezh2*;*Ai9* WT and KO mice showing decrease in Tom+ cells in the KO mouse. Scale bar = 100 μm. **B.** Graph displaying the density of Tom+ cells in P5 cortex. All stats are unpaired t-tests: *** = p ≤ .001, ** = p ≤ .01. n = 4 WT and 4 KO brains, from 4 different litters.

### Changes in gene expression and chromatin accessibility in the MGE of *Ezh2* KO mice

We then investigated whether transcriptomic and epigenetic changes in the *Ezh2* KO MGE are predictive of these cell fate changes in the mature forebrain. We generated single nuclei suspensions from E12.5 and E15.5 MGE and used the 10x Genomics Multiome kit to define the gene expression profile and chromatin accessibility within individual cells. We obtained a total of 51,656 nuclei that passed QC (Supplementary Fig. 6A), with the following breakdown of cells per age and genotype: E12.5 WT = 6,391; E12.5 Het = 6,608; E12.5 KO = 8,546; E15.5 WT = 11,477; E15.5 Het = 10,027; E15.5 KO = 8,607. These nuclei can be clustered based on their transcriptome, chromatin accessibility or integrated RNA and ATAC using the Weighted Nearest Neighbor (WNN) function in Seurat^72^ (Fig. 7A and Supplementary Fig. 6B). Analysis of several genes revealed the expected progression from radial glia cells (*Nes*) to immature neuronal progenitors (*Ascl1*) to postmitotic neural precursors (*Dcx* and *Rbfox3*), which was verified with pseudotime analysis (Fig. 7B).

**Fig. 7.**
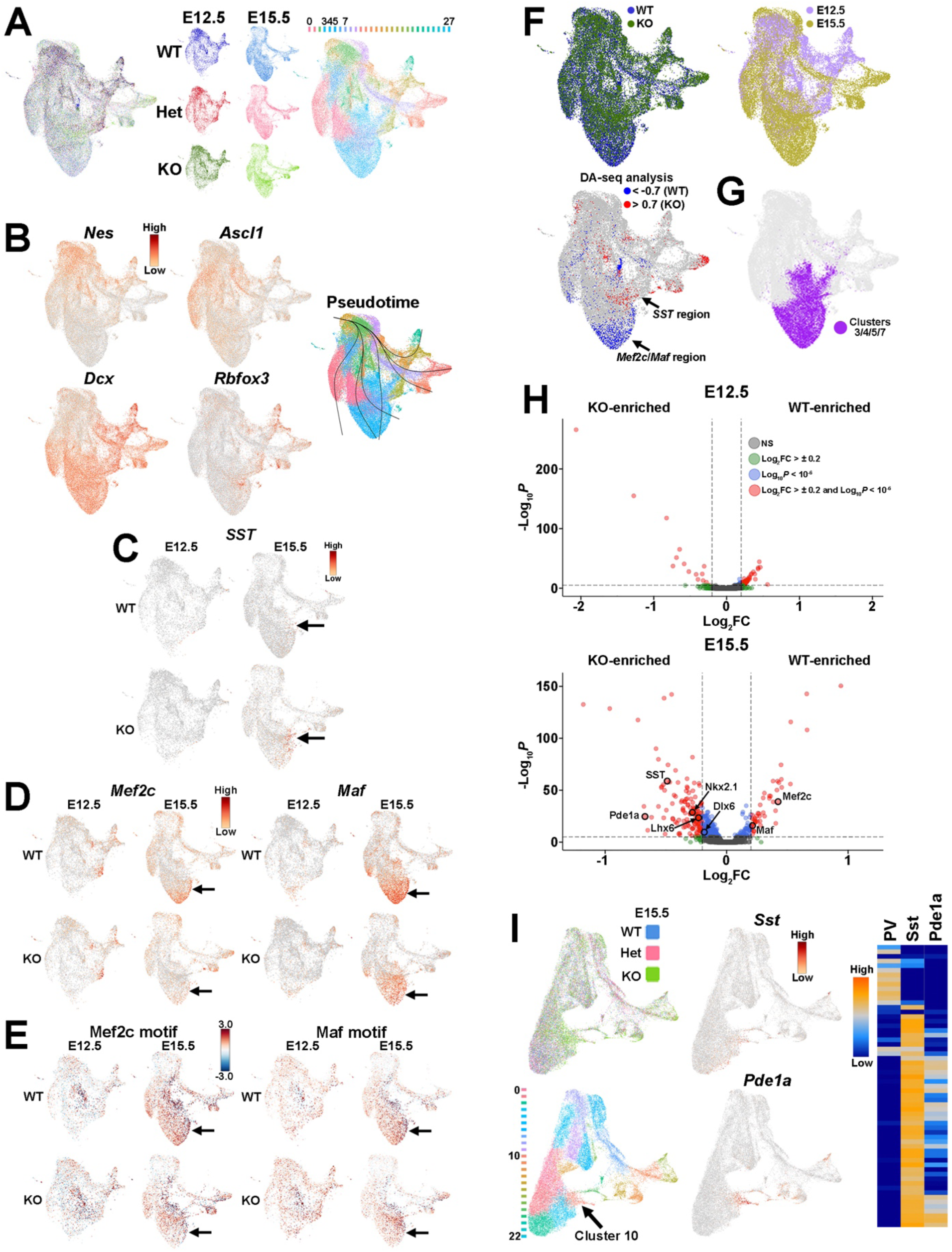
Shifts in transcriptome and differential abundance cell types in *Ezh2* KO mice. **A.** Uniform Manifold Approximation and Projection (UMAP) plots of E12.5 and E15.5 integrated single cell RNA- and ATAC-seq (Multiome) dataset via weighted nearest neighbor (WNN), annotated by age and genotype (left and middle) or putative cell clusters (right). Labels for putative cell clusters are listed above the UMAP. **B.** Markers for radial glia cells (*Nes*), cycling GE progenitors (*Ascl1*) and postmitotic immature neurons (*Dcx*, *Rbfox3*), with general trajectory confirmed by pseudotime. **C.** *SST* is enriched in E15.5 KO MGE postmitotic neurons compared WT MGE. **D.** Enrichment of *Mef2c* and *Maf*, two genes predictive of PV-fated interneurons, in E15.5 WT MGE compared to KO. **E.** Enrichment of *Mef2c* and *Maf* binding motifs in E15.5 WT MGE compared to KO. **F.** E15.5 integrated RNA and ATAC dataset annotated by genotype (left) and age (right). **G.** Differential abundance (DA) analysis using DA-seq reveals that *SST*+ cells are located in KO-enriched clusters (red) whereas *Mef2c*- and *Maf*-expressing cells are located in WT-enriched clusters (blue). DA score threshold of +/- 0.7 used for significant enrichment of DA cells in DA-seq analysis. **H.** Clusters 3, 4, 5 and 7 from panel **A** that are enriched for PV- and SST-fated cells. **I.** Volcano plots depicting genes enriched in WT or KO MGE at E12.5 (top) and E15.5 (bottom). Thresholds used were Log_2_ fold change (FC) > ± 0.2 and a false discovery rate (FDR) of Log_10_P < 10^-6^. **J.** E15.5 integrated RNA and ATAC dataset annotated by genotype (top left) and putative cell clusters (bottom left). The top gene enriched in the cluster harboring *SST*+ cells (Cluster 10, arrow) is *Pde1a*. *Pde1a* is expressed in many SST+ interneuron subtypes in the adult mouse (each row is a mature interneuron subtype), but excluded from PV+ interneurons (right, adapted from the Allen Brain Map Transcriptomics Explorer).

Since there was a general increase in SST+ interneurons in the *Ezh2* KO mouse, we examined *SST* expression in this single cell dataset. While no obvious differences in *SST* expression was apparent at E12.5, we did observe an increase in *SST* expression in the MGE of E15.5 KO compared to WT (Fig. 7C). While PV is not expressed in the embryonic mouse brain, two genes that are enriched in PV-fated interneurons and critical for their development are the transcription factors *Mef2c* and *Maf*^21,73^. We found that both genes are strongly reduced in E15.5 KO MGEs compared to WT (Fig. 7D). Complementing this gene expression analysis, the motifs for these transcription factors are enriched in accessible regions of the E15.5 WT MGE compared to KO mice (Fig. 7E). Thus, our gene expression analysis reveals an apparent increase in *SST* expression and decrease in *Maf* and *Mef2c* expression in the MGE of E15.5 KO mice, which is consistent with the increase in SST+ and decrease of PV+ interneurons in *Ezh2* KO brains.

Additionally, WT and KO cells appeared to display differential abundance (DA) in specific regions of the UMAP plot, most notably in the E15.5 dataset (Fig. 7F and Supplementary Fig. 6B). To confirm this observation, we performed DA analysis using DA-seq^74^. DA-seq determines a DA score for each cell, whereby a cell that is surrounded by KO cells in a k-nearest neighbor (KNN) graph has a score closer to +1, and a cell surrounded by WT cells has a score closer to -1. This DA score does not rely upon previously identified clusters, and it does not require similar cell numbers between different conditions. Our DA-seq analysis revealed that most cells with a DA score above +0.7 or below -0.7 were from the E15.5 MGE (Fig. 7G), whereas cells from the E12.5 MGE displayed little differential abundance. Furthermore, most cells with a DA score below -0.7 (blue, WT bias) were in the clusters enriched for *Maf* and *Mef2c*, which are putative PV+ interneurons (Fig. 7G). The cluster containing *SST*+ cells contained numerous cells with DA score above 0.7 (red, KO bias) (Fig. 7G). Thus, we observe a differential abundance of WT and KO cells specifically in the E15.5 dataset, with an increase of KO cells in clusters expressing *SST* and an increase of WT cells in clusters where *Maf* and *Mef2c* are strongly enriched.

To specifically focus on these clusters enriched for SST- and PV-fated interneurons, we extracted the four clusters containing these cells from the dataset (clusters 3, 4, 5 and 7) (Fig. 7A,H). Using thresholds of a Log_2_ fold change (FC) > ± 0.2 and a false discovery rate (FDR) of Log_10_P < 10^-6^, we identified 59 differentially expressed genes at E12.5 (46 downregulated and 15 upregulated in KO) and 176 differentially expressed genes at E15.5 (46 downregulated and 130 upregulated in KO) (Fig. 7I and Supplementary Tables 1-2). Notably, both *Maf* and *Mef2c* were significantly enriched in the E15.5 WT MGE whereas *SST* was upregulated in the KO MGE, consistent with the observations above (Fig. 7I). Additionally, the MGE-specific transcription factors *Nkx2.1* and *Lhx6* were also upregulated in the KO MGE in these clusters.

Since differentially expressed genes of interest were restricted to E15.5, we reexamined the integrated E15.5 RNA+ATAC dataset alone (Supplementary Fig. 6B). In this dataset, *SST* was strongly enriched in cluster 10 (Fig. 7J). The top gene expressed in this cluster was Phosphodiesterase 1A, *Pde1a*, which is also significantly upregulated in the E15.5 KO MGE (Fig. 7I). Based on the Allen Brain Institute’s single cell transcriptomic adult mouse brain dataset, *Pde1a* is enriched in many SST+ interneuron subtypes while it’s excluded from PV+ interneurons (Fig. 7J), providing additional evidence that there is an increase in SST-fated cells expressing in the MGE of KO mouse. In sum, this analysis indicates that the shift in MGE-derived interneuron subtypes is most evident in the E15.5 MGE, with an increase in both *SST* expression levels and the number of *SST*-expressing cells in the MGE of *Ezh2* KO mice, with a corresponding decrease in PV-fated cells and PV-predictive genes.

### Changes of H3K27me3 at distinct genomic loci in MGE of *Ezh2* KO mice

H3K27me3 levels are strongly downregulated in the MGE of *Ezh2* KO mice (Fig. 1). To look at H3K27me3 changes at specific genes, we performed bulk CUT&Tag^75^ with a H3K27me3 antibody in the MGE of WT and KO mice. We did not normalize total reads to a spike-in or *E. coli* DNA control so that the global downregulation of H3K27me3 in the KO was intentionally ignored from this analysis and instead we could focus on the relative changes of H3K27me3 levels at specific loci between genotypes. We performed 3 biological replicates for each age and genotype, using Epic2^76^ for peak calls and the DiffBind package^77^ with edgeR’s trimmed mean of M values (TMM)^78^ for comparative analysis. Biological replicates were grouped together in both principal component analysis (PCA) and unbiased correlation heatmaps, with age being a greater differentiation factor compared to genotype (Fig. 8A-B).

**Fig. 8.**
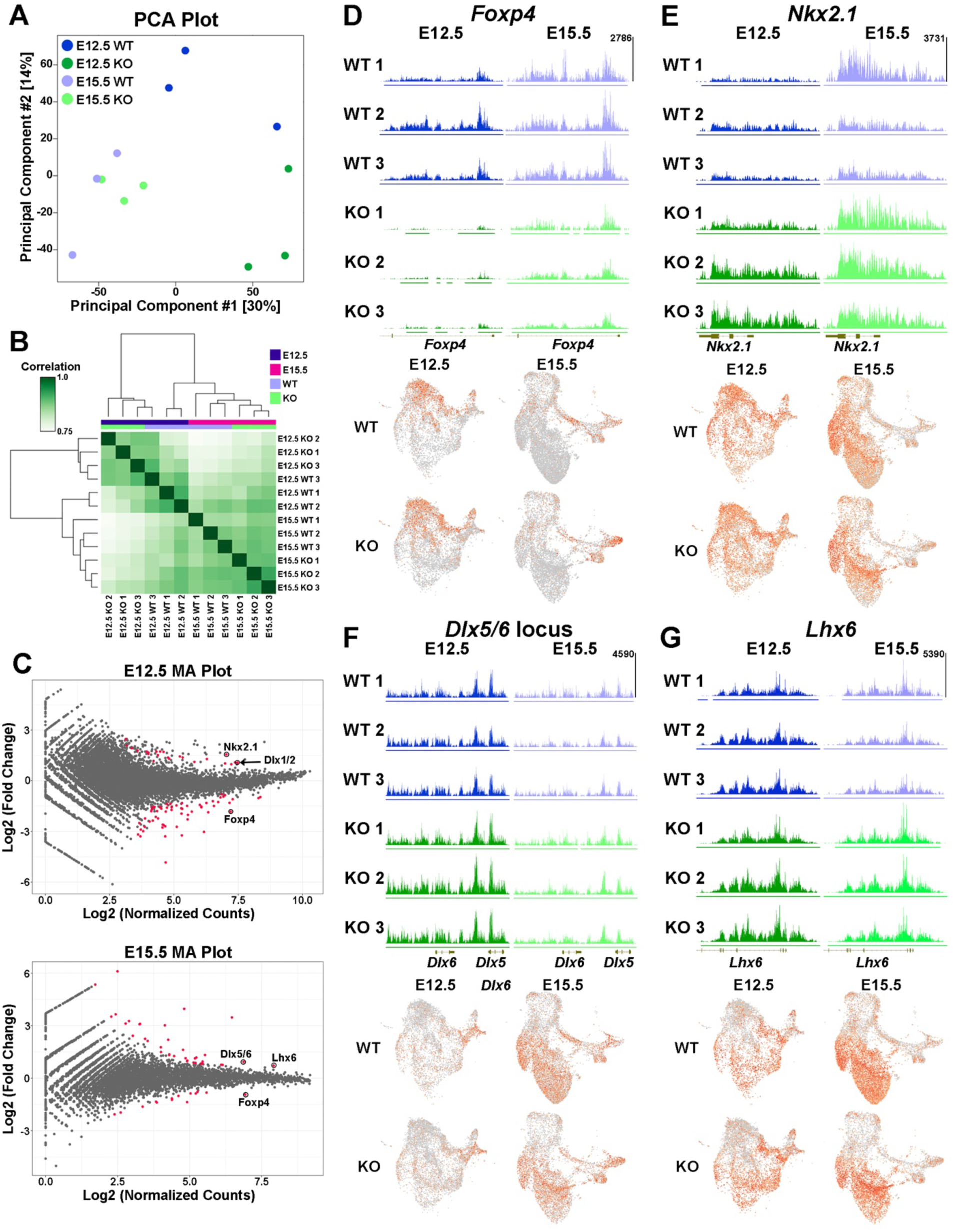
Altered H3K27me3 at distinct genomic loci in *Ezh2* KO MGE. **A.** Principal Components Analysis (PCA) plot comparing the E12.5 WT, E12.5 KO, E15.5 WT and E15.5 KO samples, 3 biological replicates each. **B.** Unbiased hierarchical correlation heatmap comparing differential peaks between all 12 samples. **C.** MA plots depicting the fold changes vs. mean peak counts for E12.5 (top) and E15.5 (bottom) data, with all significant peaks (FDR ≤ 0.1) highlighted in red. **D-G.** H3K27me3 levels at the *FoxP4* (D), *Nkx2.1* (E), *Dlx5/6* (F) and *Lhx6* (G) loci in all 12 samples (top) along with the integrated UMAP plots showing gene expression profiles in the four different conditions (bottom).

Comparative analyses revealed significant changes in the relative levels of H3K27me3 at 82 loci at E12.5 (59 decreased and 23 increased loci in KO) and 51 loci at E15.5 (14 decreased and 37 increased loci in KO) (Fig. 8C and Supplementary Tables 3-4). One of the most prominent changes was a strong reduction of H3K27me3 at the *Foxp4* locus at both E12.5 and E15.5 (Fig. 8D). Despite the strong decrease in H3K27me3 signal, we do not observe a concomitant increase in *Foxp4* gene expression in the KO MGE (Fig. 8D). While *Foxp4* is strongly enriched in the LGE^79^, its function in forebrain development has not been explored. Surprisingly, several genes regulating development of MGE-derived interneurons displayed a relative increase in H3K27me3 levels in the KO MGE: *Nkx2.1* and the *Dlx1/2* locus were increased at E12.5 whereas *Lhx6* and the *Dlx5/6* locus were increased at E15.5 (Fig. 8C, E-G). Like *Foxp4*, we did not observe a corresponding decrease in expression levels of these genes in the KO MGE. In fact, several of these genes displayed increased expression in the postmitotic SST- and PV-fated cell clusters (Fig. 7I).

Thus, even though there is a strong global decrease of H3K27me3 in the *Ezh2* KO MGE, we identified differential changes in the relative level of H3K27me3 at various loci in the genome. This indicates that specific genomic loci are more susceptible (e.g., *Foxp4*) or resistant (e.g., *Nkx2.1*) to H3K27me3 loss in the absence of *Ezh2*. The mechanism by which loss of *Ezh2* generates these differential effects at genomic loci, and how these changes in H3K27me3 levels relate to gene expression, require further investigation.

## DISCUSSION

There is growing evidence that dysregulation of epigenetic mechanisms can lead to a variety of human diseases and neurodevelopmental disorders^36–38,80,81^. For example, postmortem tissue from schizophrenic patients displays alterations in genome organization and other epigenomic characteristics^82,83^. Additionally, many genes associated with neurological and psychiatric diseases are enriched in immature interneurons during embryonic development^9,10,84,85^. Thus, advancing our knowledge of gene regulation mechanisms during interneuron development is critical for understanding both normal development and disease etiologies.

In this study, we find that loss of *Ezh2* in the MGE decreases the density and proportion of PV+ cells, often with a corresponding increase in SST+ cells. This shift in interneuron fate was most prominent in the cortex and the CA2/3 region of the hippocampus, with an overall decrease in total MGE-derived interneurons also observed in the cortex. A decrease in PV+ cells was observed in the CA1, DG and striatum without a significant increase in SST+ cells (Figs. 2-3 & Supplementary Fig. 1-3). In the hippocampus, we also observe an increase in MGE-derived nNos+ cells in the KO (Fig. 3). Unlike PV+ and SST+ interneurons, the spatial and temporal origin of hippocampal nNos+ in the MGE is not well characterized, so it’s unclear how loss of *Ezh2* increased nNos+ cells. These phenotypes were due to *Ezh2* function in cycling MGE progenitors, as no changes were observed when *Ezh2* was removed in postmitotic MGE cells (Supplementary Fig. 5). This differential severity of interneuron fate changes in specific brain regions highlights the importance of examining interneurons in multiple brain regions.

In particular, CA2/3 displayed a significant shift in interneuron fate compared to other hippocampal regions, more closely matching changes in the cortex. CA1 and CA2/3 have similar densities and percentages of MGE-derived interneurons (Fig. 3 & Supplementary Fig. 3). Why these regions display such different phenotypes in the KO remains unknown. The severity of CA2/3-specific changes could arise from differences in interneuron migration, cell survivability or specific circuit interactions within this region. Additionally, the high density of MGE-derived oligodendrocytes in the WT CA2/3 was quite striking, as we did not observe these cells in the cortex or other hippocampal regions. MGE-derived oligodendrocytes do migrate into the cortex, but they are almost entirely eliminated during the second postnatal week^86^. We are unaware of any reports describing MGE-derived oligodendrocytes in the hippocampus, and thus further studies are needed to understand why a population of MGE-derived oligodendrocytes perdures in CA2/3 into adulthood. Assuming these MGE-derived oligodendrocytes are generated towards the end of the cell cycle, then loss of this population is consistent with preemptive cell cycle exit in the *Ezh2* KO mouse and the overall reduced production of MGE-derived cortical interneurons observed at P5 (Fig. 6).

Despite these changes in cell fate, mature PV+ and SST+ cortical interneurons displayed normal intrinsic physiological properties. However, cortical PV+ interneurons displayed significantly greater axon length and complexity in the KO mouse. This could be a form of compensation, as one way to increase inhibition with fewer PV+ cells is for the surviving PV+ interneurons to have increased synaptic contacts. Alternatively, enhanced activation of PV+ interneurons can result in more elaborate axon morphologies^87^: if individual PV+ interneurons are receiving more glutamatergic inputs because of decreased number of PV+ cells in *Ezh2* KO mice, than this could in part explain the morphological changes. Additionally, PV+ interneurons normally form exuberant axon projections and synaptic connections during the first two postnatal weeks which are then pruned and retracted during the 3^rd^ and 4^th^ weeks^88^. Thus, it’s possible that this normal developmental retraction is disrupted in the KO mouse due to fewer PV+ cortical interneurons.

Single cell sequencing revealed that these cell fate changes are already apparent in the developing MGE. First, the majority of transcriptome differences were observed in the E15.5 MGE while the E12.5 MGE from WT and KO mice were quite similar. This could imply that the loss of *Ezh2* takes time to manifest (due to perdurance of H3K27me3 marks, rate of histone turnover, etc.), with the strongest affects becoming more evident towards the end of MGE neurogenesis. Second, there was an increase in *SST* expression and a decrease in expression of PV-predictive genes *Mef2c* and *Maf* in the *Ezh2* KO mouse at E15.5 (Fig. 7C-D,I). Third, differential abundance analysis reveals that there is a strong bias for cells from the KO MGE in the *SST*-expressing region, and an enrichment for WT MGE cells in the *Mef2c*/*Maf*-expressing region (Fig. 7F). Fourth, the top gene expressed in the *SST*-enriched cluster at E15.5 is *Pde1a*, which is expressed by many mature *SST*+ interneuron subtypes but excluded from *PV*+ interneurons (Fig. 7J). In sum, these transcriptional and cellular differences in the MGE are likely determinative for the shifts in SST+ and PV+ interneurons in the adult brain of *Ezh2* KO mice.

Despite the global downregulation of H3K27me3, we found that loss of *Ezh2* had differential effects on relative H3K27me3 levels at specific gene loci. One of the loci most susceptible to *Ezh2* loss was the *Foxp4* locus, with a significant loss of H3K27me3 in the KO MGE at E12.5 and E15.5 (Fig. 8C-D). Surprisingly, a significant increase in *Foxp4* expression (predicted based on H3K27me3 downregulation) was not observed. *Foxp4* is enriched in the LGE^79^, but it has not been well-studied in neurodevelopment. In heterologous cell lines, FOXP4 can directly interact with the transcription factors SATB1, NR2F1 and NR2F2^89^, all of which are critical for development of MGE-derived interneurons^90,91^. Whether these interactions occur in the developing brain is unclear. A study on medulloblastoma found that *Foxp4* and *Ezh2* are both targets of the microRNA miR-101-3p^92^, indicating the function of these genes might be linked in some scenarios. Why the *Foxp4* locus is extremely sensitive to *Ezh2* loss requires further study.

In the *Ehz2* KO MGE, we observed a significant relative increase in H3K27me3 levels at several transcription factors critical for development of MGE-derived interneurons: *Nkx2.1* and *Dlx1/2* locus at E12.5 and *Lhx6* and *Dlx5/6* locus at E15.5. This raises the possibility that some genes playing critical roles in fate determination may be more resistant to epigenetic changes, in this case loss of *Ezh2*. For example, the interaction between *Sox2*, a transcription factor essential in the epiblast of pre-implantation embryos, and a critical enhancer downstream is maintained even when artificial boundaries are introduced between these regions^93^. As *Nkx2.1* is a ‘master regulator’ of MGE fate, and the *Nkx2.1* locus displays unique chromatin organization in the MGE^23^, it could be more resistant to epigenetic modifications.

Similar to *Foxp4*, we did not observe a corresponding change in the global expression of these transcription factors that is predictive of these H3K27me3 changes. In fact, we actually observed a significant increase in *Nkx2.1* and *Lhx6* expression in the postmitotic SST- and PV-fated clusters (Fig. 7I), which is in contrast to the predicted decreased expression based on H3K27me3 levels. While our previous results showed a strong relationship between H3K27me3 and gene repression in the embryonic mouse brain^23^, the correlation here is weaker. It will be interesting to explore gene-specific changes in the resistance or susceptibility to epigenetic changes going forward, both in terms of normal development and regarding neurodevelopmental and psychiatric diseases.

## METHODS

### Animals

All experimental procedures were conducted in accordance with the National Institutes of Health guidelines and were approved by the NICHD Animal Care and Use Committee (protocol #20-047). The following mouse lines were used in this study: *Nkx2.1-Cre* (Jax# 008661)^60^, *Ezh2^F/F^* (Jax# 022616)^94^, *Dlx5/6-Cre* (Jax# 008199)^95^ and *Ai9* (Jax# 007909)^96^. For timed matings, noon on the day a vaginal plug was observed was denoted E0.5. Both embryonic and adult male and female embryonic mice were used without bias for all experiments.

### Harvesting and Fixing brain tissue

#### MGE dissections

E12.5 and E15.5 embryos were removed and placed in ice-cold carbogenated ACSF. Tails were clipped for PCR genotyping. During genotyping, embryonic brains were harvested and MGEs were dissected from each embryo and stored in ice-cold carbogenated ACSF. For E12.5, the entire MGE was removed as previously described^97^. For E15.5, the MGE was dissected under a fluorescent dissecting microscope to ensure collection of only Tom+ MGE cells and minimize collection of postmitotic MGE-derived cells in the striatum anlage and other ventral forebrain structures. Upon obtaining genotyping results (∼90 minutes), MGEs from embryos of the same genotype were combined (when applicable) to generate single nuclei dissociations.

#### Postnatal brain fixations

All mice ≥ P5 were terminally anesthetized with an i.p. injection of Euthasol (270 mg/kg, 50 μl injection per 30 g mouse) and perfused with 4% paraformaldehyde (PFA). In some cases, tail snips were collected for genotyping prior to perfusion. Brains were removed and post-fixed in 4% PFA O/N at 4°C, then cryoprotected in 30% sucrose in PBS O/N at 4°C before embedding in OCT. Brains were sectioned at 30 mm on a CryoStar^TM^ NX50 cryostat and stored as floating sections in antifreeze solution (30% ethylene glycol, 30% glycerol, 40% PBS) at -20°C in 96-well plates.

#### Embryonic brain fixations

Pregnant dams were terminally anesthetized with an i.p. injection of Euthasol (270 mg/kg, 50 μl injection per 30 g mouse). E12.5-E15.5 embryos were removed and placed in ice cold carbogenated artificial cerebral spinal fluid (ACSF, in mM: 87 NaCl, 26 NaHCO_3_, 2.5 KCl, 1.25 NaH_2_PO_4_, 0.5 CaCl_2_, 7 MgCl_2_, 10 glucose, 75 sucrose, saturated with 95% O_2_, 5% CO_2_, pH 7.4). Embryonic brains were removed and incubated in 4% PFA O/N at 4°C. In some cases, tail snips were collected for genotyping. Brains were washed in PBS, transferred to 30% sucrose in PBS and embedded in OCT upon sinking. Brains were sectioned at 14-16 μm, mounted directly onto Permafrost slides, and stored at -80°C.

### Immunohistochemistry & Fluorescent *In Situ* Hybridizations (ISH)

30 μm free floating sections from P30-40 brains were washed in PBS to remove antifreeze solution and incubated in blocking solution (10% Normal Donkey Serum in PBS + 0.3% Triton X-100) for 1-2 hours. Sections were incubated with primary antibodies in blocking solution for 48 hours at 4°C, then washed in PBS for 2-4 hours at RT. Sections were incubated with secondary antibodies with DAPI in blocking solution O/N at 4°C, washed in PBS, mounted and imaged.

Cryosectioned E12.5-E15.5 brains sections were incubated with blocking buffer (10% Normal Donkey Serum in PBS + 0.1% Triton X-100) for 1 hr at RT, then incubated in primary antibody solution in blocking buffer O/N at 4°C, washed in PBS for 1-2 hrs, then incubated with secondary antibodies with DAPI in blocking solution for 2 hrs at RT or O/N at 4°C. Slides were washed in PBS and imaged. The following antibodies were used in this study: rabbit-anti H3K27me3 (1:100, Cell Signaling 9733T), rat-anti SST (1:300, Millipore MAB354), goat-anti PV (1:1000, Swant PVG213), rabbit-anti PV (1:1000, Swant PV27), rabbit-anti nNos (1:500, Millipore MAB5380), rabbit anti-Olig2 (1;500, Millipore AB9610). Species-specific fluorescent secondary antibodies used were conjugated to AlexaFlour^®^ 488, 647 and 790, and all used at 1:500.

RNAscope^TM^ ISH assays (Advanced Cell Diagnostics) were performed on E12.5-E13.5 brain sections according to the manufacturer’s instructions. The following probes were used in this study: *Nkx2.1* (434721), *Ezh2* (802751-C3) and *tdTomato* (317041-C2).

All images were captured at 20X on a Zeiss Axioimager.M2 with ApoTom.2 (with Zen Blue software) or an Olympus VS200 Slide Scanner (VS200 ASW). Image post-processing was performed with Adobe Photoshop and ImageJ software.

### Western Blots and Analysis

Core histone proteins from E13.5 MGE samples (pooled from 2-4 brains per genotype) were extracted using EpiQuik Total Histone Extraction kit (Epigentek# OP-0006). We obtained 40-80 μg of histone proteins per extraction and loaded 20 μg of protein onto a 4-12% Bis-Tris Plus mini gel (Invitrogen# NW04120BOX). Gels were run for ∼20 minutes at 200V using the Invitrogen mini gel tank with Blot MES SDS Running buffer. Gels were transferred to iBlot2 polyvinylidene difluoride (PVDF) membranes (Invitrogen# IB24001) in the iblot2 at 20V for 7 minutes. Blots were incubated with the primary antibody, mouse anti-H3 (Cell Signaling Technology# 3638S; 1:1000) and Rabbit anti-H3K27me3 (Cell Signaling Technology# 9733S;1:500) overnight at 4°C, and then incubated with secondary anti-mouse-Starbright Blue-520 (Bio-Rad# 64456855; 1:2000) and anti-Rabbit-Starbright Blue-700 (Bio-Rad# 64484700; 1:2000) for 1 hour at room temperature. Blots were imaged on a Bio-Rad ChemiDoC MP imaging system.

For analysis, a box was drawn around the ROI in each lane, with the same sized box used for both H3 and H3K27me3 signals for the WT, Het and KO lanes in each gel. The average (mean) gray value was calculated in each box, then a lane-specific background signal taken just below each ROI was subtracted from each value. For normalization, each H3K27me3 value was divided by the corresponding H3 value for each lane (e.g., WT H3K27me3/WT H3). Then these Het and KO values were divided by the WT value to determine the % of H3K27me3 signal compared to WT.

### Cell Counting

#### Adult brains

All cell counts were performed by hand and blind to genotypes. Total brains counted for *Nkx2.1-Cre;Ezh2;Ai9* mice: WT = 5, Het = 5, KO = 6, from 4 different litters. Total brains counted for *Dlx5/6-Cre;Ezh2;Ai9* mice: WT = 4, Het = 3, KO = 5, from 3 different litters. Counted cells consisted of either Tom+, Tom+/SST+, or Tom+/PV+ (and in the hippocampus, Tom+/nNos+); any SST+, PV+ or nNos+ cells that were Tom-were excluded from counts since they likely did not recombine at the *Ezh2* (or Ai9) locus. For all sections, area was calculated using ‘Measurement’ function in Photoshop, and all average areas described below include WT, het and KO brains. Cortex: Counts were performed in somatosensory cortex on 3 non-consecutive sections per brain, then averaged together. Individual cortical images were divided into superficial and deep sections using DAPI staining to define the layer III-IV boundary. Average cortical area/section = 0.85 mm^2^. Striatum: Counts were performed on 3 sections per brain, one section each through the anterior, middle and posterior striatum, then averaged together. Average striatal area/section = 3.15 mm^2^. Hippocampus: Counts were performed on 8 non-consecutive sections per brain, then averaged together. More hippocampal sections were counted per brain due to the comparatively low number and section-to-section variability of Tom+ cells in the hippocampus. Sections were restricted to the anterior and middle hippocampus; the posterior hippocampus was excluded due to greater variability in interneuron density in this region. Hippocampal sections were divided into CA1, CA2/3 and DG regions using DAPI staining (Fig. 3A). Small Tom+ cell bodies (identified as oligodendrocytes) in CA2/3 (Fig. 4) were excluded from interneuron counts and counted as separate group. Average CA1 area/section = 0.98 mm^2^, average CA2/3 area/section = 0.49 mm^2^, average DG area/section = 0.60 mm^2^.

#### P5 brains

All cell counts were performed by hand and blind to genotypes. Total brains counted for *Nkx2.1-Cre;Ezh2;Ai9* mice: WT = 4 and KO = 4, from 4 different litters. Counts were performed in somatosensory cortex on 3 non-consecutive sections per brain, then averaged together. Individual cortical images were divided into superficial and deep sections using DAPI staining. Average cortical area/section = 0.92 mm^2^.

### *In vitro* electrophysiology

#### Slice preparation

Mice were anesthetized with isoflurane (5% isoflurane (vol/vol) in 100% oxygen), perfused transcardially with an ice-cold sucrose solution containing (in mM) 75 sucrose, 87 NaCl, 2.5 KCl, 26 NaHCO_3_, 1.25 NaH_2_PO_4_, 10 glucose, 0.5 CaCl_2_, and 2 MgSO_4_, saturated with 95% O_2_ and 5% CO_2_ and decapitated. Brain was rapidly removed from the skull and transferred to a bath of ice-cold sucrose solution. Coronal slices of 300 µm were made using a vibratome (Leica Biosystems) and were stored in the same solution at 35°C for 30 min and at room temperature (RT) for an additional 30-45 min before recording.

#### Electrophysiology

Whole-cell patch clamp recordings on tdTomato+ cells in cortical layers V/VI cells were performed in oxygenated ACSF containing (in mM) 125 NaCl, 2.5 KCl, 26 NaHCO_3_, 1.25 NaH_2_PO_4_, 10 glucose, 2 CaCl_2_ and 1 MgCl_2_. The ACSF was equilibrated with 95% O_2_ and 5% CO_2_ throughout an entire recording session which typically lasted between 30 minutes to 1 hour to ensure sufficient permeation of neurobiotin. Recordings were performed at 30°C-33°C. Electrodes (3-7 MΩ) were pulled from borosilicate glass capillary (1.5 mm OD). The pipette intracellular solution contained (in mM) 130 potassium gluconate, 6.3 KCl, 0.5 EGTA, 10 HEPES, 5 sodium phosphocreatine, 4 Mg-ATP, 0.3 Na-GTP and 0.3% neurobiotin (pH 7.4 with KOH, 280-290 mOsm). Membrane potentials were not corrected for the liquid junction potential. During patching, cell-attached seal resistances were >1 GΩ. Once whole-cell configuration was achieved, uncompensated series resistance was usually 5-30 MΩ and only cells with stable series resistance (<20% change throughout the recording) were used for analysis. Data were collected using a Multiclamp 700B amplifier (Molecular Devices), low-pass filtered at 10 kHz and digitally sampled at 20 kHz, and analyzed with pClamp10 (Molecular Devices). To characterize the intrinsic membrane properties of neurons, hyperpolarizing and depolarizing current steps were injected at 0.1 Hz under current-clamp configuration.

#### Data analysis

All intrinsic properties were measured in current-clamp configuration and calculated from 800 millisecond-long current injections unless noted otherwise. The resting membrane potential (in mV) was measured with 0 pA current injection a few minutes after entering whole-cell configuration. All other properties were measured holding the cell at -70 mV. Input resistance (in MΩ) was calculated using Ohm’s law from averaged traces of 100 ms long negative current injections of -20 pA. Action potential (AP) threshold was calculated as the potential when voltage change over time was 10 mV/ms using the first observed spike. AP amplitude (in mV) was calculated as the time difference in potential from the spike peak to spike threshold. AP/spike half-width (in ms) was calculated as the difference in time between the ascending and descending phases of a putative spike at the voltage midpoint between the peak of spike and spike threshold. Adaptation ratio was calculated as the ratio of the number of APs in the number of spikes in the last 200 ms over the number of APs in the first 200 ms of a positive current injection that elicited approximately 20 Hz firing. Afterhyperpolarization (AHP) amplitude was calculated as the difference between AP threshold and the most negative membrane potential after the AP, measured on the response to the smallest current step evoking an AP (Rheobase). Membrane time constant (in ms) was determined from a monoexponential curve best fitting the falling phase of the response to a small hyperpolarizing current step.

### CUBIC clearing and streptavidin staining

After performing electrophysiological recordings, brain slices were fixed in 4% PFA in 0.1M PB and kept overnight at 4°C and then kept in 20% sucrose (in PB). The brain slices were processed for CUBIC (Clear, Unobstructed Brain/Body Imaging Cocktails and Computational analysis) clearing^98^. Slices were first washed with 0.1M PB (3 times for 10 min) at RT, followed by immersion in CUBIC reagent 1 for two days at 4°C. After two days of incubation, slices were washed with 0.1M PB (4 times for 30 min) at RT to ensure complete removal of CUBIC reagent 1. Slices were then incubated in fluorophore-conjugated streptavidin (1:500; ThermoFisher Scientific) in 0.1M PB (0.5% TritonX-100) overnight at 4°C. Slices were subsequently washed with 0.1M PB (4 times, 30 min) at RT and mounted with CUBIC reagent 1. Filled neurons were imaged with a Nikon A1R confocal microscope. Z-stacked images (each stack 1 µm) were acquired with a 40X oil-immersion objective.

### Sholl analysis

Neurobiotin-filled neurons were processed and reconstructed using the Neurolucida 360 software (NL360) (MBF Biosciences). Briefly, image stack files were converted to JPEG 2000 file format with a MicroFile+ software, and the converted images were loaded into the NL360 software package. Neurites of PV were then traced manually in a 2D environment. Among the traced neurites, dendrites were easily distinguishable from axons, whose extensive ramifications maintained a constant diameter and had varicosities. Branched structure and Sholl analysis were performed using the built-in functions of the Neurolucida explorer, in which a series of concentric spheres (10 µm interval between radii) were created around the middle point (soma of the traced neuron).

### Generating Single Nuclei Suspensions for CUT&Tag and Multiome Experiments

Single nuclei suspensions were prepared as previously describes^22,97^ with slight modifications. *CUT&Tag*: MGEs were transferred to a 1 mL Dounce homogenizer containing DNA Lysis Buffer (10 mM Tris-HCl (Ph.7.4), 10 mM NaCl, 3 mM CaCl_2_, 0.1% Tween-20, 1.5% BSA and 0.1% IGEPAL CA-630 in nuclease-free water, 1 mL per sample). Cells were dounced with pestle A and pestle B, ∼10-15 times each, and pipetted through a 40 μm filter onto a pre-chilled 50 mL conical tube on ice and wet with 1 mL of DNA Nuclei Wash Buffer (in mM: 10 Tris-HCl (Ph.7.4), 10 NaCl, 3 CaCl_2_, with 0.1% Tween-20 and 1.5% BSA in nuclease-free water, 5 mL per sample). Transfer lysed nuclei suspension through pre-wetted filter, then rinse dounce with 1 mL Nuclei Wash Buffer and transfer through filter. Nuclei suspension was divided into 2 pre-chilled 2 mL tubes and spun at 500 g for 5 minutes at 4°C. After removing supernatant, nuclei pellet was washed once with 1 mL Nuclei Wash Buffer and then once with 1 mL of 1X CUT&Tag Wash Buffer (from Active Motif CUT&Tag IT Assay kit). Remove CUT&Tag Wash Buffer, leaving ∼30-50 μL in each, triturate and combine nuclei suspensions from both tubes. Nuclei concentration was determined on Countess II FL Automated Cell Counter. 100,000-125,000 nuclei were used for each CUT&Tag reaction using the CUT&Tag IT Assay Kit (Active Motif, #53610) per manufacturer’s instructions. Primary antibody was rabbit-anti H3K27me3 (Cell Signaling, 9733T, 1:50) or rabbit-anti H3K27ac (Cell Signaling XXX; 1:50), and secondary antibody was guinea pig anti-rabbit (Active Motif, 105465 from CUT&TagIT kit, 1:100).

#### 10x Genomics Multiome kit

MGEs were transferred to a 1 mL Dounce homogenizer containing Multiome Lysis Buffer (in mM: 10 Tris-HCl (Ph.7.4), 10 NaCl, 3 CaCl_2_, 1 DTT, with 0.1% Tween-20, 1.5% BSA and 0.1% IGEPAL CA-630 in nuclease-free water, 1 mL per sample). Lyse cells by douncing with pestle A and pestle B, ∼10-15 times each. Place a 40 μm filter onto a pre-chilled 50 mL conical tube on ice and wet with 1 mL of Multiome Nuclei Wash Buffer (in mM: 10 Tris-HCl (Ph.7.4), 10 NaCl, 3 CaCl_2_, 1 DTT, with 0.1% Tween-20, 1.5% BSA and 1 U/μL in nuclease-free water, 5 mL per sample). Transfer lysed nuclei suspension through pre-wetted filter, then rinse dounce with 1 mL Multiome Wash Buffer and transfer through filter. Divide nuclei suspension into 2 pre-chilled 2 mL tubes and spin at 500 g for 5 minutes at 4°C. After removing supernatant, wash nuclei pellet twice with 1 mL Multiome Wash Buffer and spin as above. Remove Multiome Wash Buffer, leaving ∼20-30 μL of solution in each tube. Triturate solution to dissociate pellets, then combine nuclei suspensions into 1 tube. Nuclei concentration was determined on Countess II FL Automated Cell Counter. Nuclei suspensions were diluted to ∼3,000-4,000 nuclei/μL, with 5 μL being used for 10x Genomics Multiome kit per manufacturer’s instructions. E15.5 data: MGE from 4 WT, Het and KO embryos from 2 different E15.5 litters were combined to generate 1 biological rep. E12.5 data: MGE from 1 WT, Het and KO mouse from a single E12.5 litter were used to generate 1 biological rep. Total cell numbers that passed QC: E12.5 WT = 6,391; E12.5 Het = 6,608; E12.5 KO = 8,546; E15.5 WT = 11,477; E15.5 Het = 10,027; E15.5 KO = 8,607.

### CUT&Tag Sequencing & Analysis

Following library amplification, DNA quantity was determined with a Qubit and library quality characterized with an Agilent Tapestation. Libraries were balanced for DNA content and pooled before performing a final SPRIselect bead 1x left size selection and paired-end sequenced (50 x 50 bp) on an Illumina NovaSeq. Sequencing read depths per library are: E12.5_WT1=12,928,439; E12.5_WT2=47,425,165; E12.5_WT3=34,733,272; E12.5_KO1=28,936,620; E12.5_KO2=27,272,500; E12.5_KO3=13,875,056; E15.5_WT1=34,525,312; E15.5_WT2=32,762,090; E15.5_WT3=39,114,390; E15.5_KO1=48,561,971; E15.5_KO2=82,783,017; E15.5_KO3: 169,597,179.

Paired-end run Illumina NovaSeq produced 2 compressed fastq.gz files for each replicate. For each age (E12.5 and E15.5) and each genotype, a total of 3 different biological reps from 3 different experiments were combined. Adaptor sequences were removed using cutadapt ^99^ v3.4 with the following parameters: -a AGATCGGAAGAGCACACGTCTGAACTCCAGTCA -A AGATCGGAAGAGCGTCGTGTAGGGAAAGAGTGT -q 20 --minimum-length 25. Trimmed reads were mapped to mouse reference genome (GRCm38/mm10) using bowtie2^100^ v2.4.2 with the following parameters: --no-unal -N 1 --no-mixed --no-discordant --very-sensitive-local –local -- phred33 -I 10. Aligned reads in sam files were further processed to remove multimappers if MAPQ was less than 10 (-q 10) and then sorted using samtools^101^ v1.12. Aligned reads that intersected blacklist regions^102^ were removed and saved to bam files using bedtools^103^ v2.30.0. PCR duplicates were removed from the bam files using picard -Xmx20g MarkDuplicates v2.25.2 (https://broadinstitute.github.io/picard/). Bigwig files were created from bam files using deepTools^104^ (v3.5.1) bamCoverage with the following parameters: --bindSize 5 --normalizeUsing RPKM. Peaks were called using Epic2^76^ on each bam file with the parameters --genome mm10 --guess-bampe and saved to bed files. Peaks were visualized using the Integrative Genomics Viewer (IGV)^105^ v2.16.1 by importing bigwig and bed files, respectively.

#### Differential analysis

Differentially enriched motifs were analyzed using edgeR^78^ v3.32.1 implemented in DiffBind^106^ v3.0.15 in R v4.0.3 (https://cran.r-project.org/). Similarity in raw peak profiling across the samples was analyzed through PCA and a sample correlation heatmap using DiffBind::plotPCA and DiffBind::plot, respectively. For differential testing, CUT&Tag reads were counted in consensus peaks with default width in DiffBind::dba.count. The counts were subsequently normalized using the TMM^107^ method by setting the normalize argument to DBA_NORM_TMM in DiffBind::dba.normalize. The false discovery rate (FDR) was determined using the Benjamini–Hochberg (BH)^108^ method for multiple hypothesis testing. Peaks with an FDR < 0.1 were considered statistically significant.

Peaks were annotated to the nearest genes using ChIPseeker package^109^ in R v4.0.3. For the TxDb and annoDb arguments in ChIPseeker::annotatePeak, we used TxDb.Mmusculus.UCSC.mm10.knownGene (DOI: 10.18129/B9.bioc.TxDb.Mmusculus.UCSC.mm10.knownGene) and org.Mm.eg.db (DOI: 10.18129/B9.bioc.org.Mm.eg.db), respectively. The transcript start site (TSS) region was defined as ranging from -5kb to +5kb. Statistically significant loci were visualized in MA plots (also known as Bland-Altman plots) using ggplot2 v3.3.3 in R.

### Multiome Sequencing & Analysis

Joint libraries for scRNA-seq and scATAC-seq were created using 10x Genomics Single Cell Multiome ATAC + Gene Expression kit (1000285) by following manufacturer’s protocols. Sequencing was conducted with paired-end (50 x 50 bp) using an Illumina HiSeq 2500 (E12.5 scATAC-seq) or NovaSeq 6000 (E12.5 scRNA-seq, E15.5 scATAC-seq, E15.5 scRNA-seq) to a following depths per library: E12.5 WT: Estimated number of cells: 7,325; Sequenced read pairs (RNA): 290,467,383; Median UMI counts per cell (RNA): 8,252; Median genes per cell (RNA): 3,281; Sequenced read pairs (ATAC): 242,612,118; Median fragments per cell (ATAC): 14,465. E12.5 Het: Estimated number of cells: 7,578; Sequenced read pairs (RNA): 198,128,926; Median UMI counts per cell (RNA): 6,006; Median genes per cell (RNA): 2,693; Sequenced read pairs (ATAC): 297,531,741; Median fragments per cell (ATAC): 16,794. E12.5 KO: Estimated number of cells: 9,989; Sequenced read pairs (RNA): 312,370,352; Median UMI counts per cell (RNA): 6,645; Median genes per cell (RNA): 2,844; Sequenced read pairs (ATAC): 210,811,269; Median fragments per cell (ATAC): 10,010. E15.5 WT: Estimated number of cells: 14,407; Sequenced read pairs (RNA): 307,184,096; Median UMI counts per cell (RNA): 5,097; Median genes per cell (RNA): 2,417; Sequenced read pairs (ATAC): 281,350,803; Median fragments per cell (ATAC): 7,808. E15.5 Het: Estimated number of cells: 12,655; Sequenced read pairs (RNA): 287,063,994; Median UMI counts per cell (RNA): 5,088; Median genes per cell (RNA): 2,401; Sequenced read pairs (ATAC): 269,896,091; Median fragments per cell (ATAC): 8,091. E15.5 KO: Estimated number of cells: 11,432; Sequenced read pairs (RNA): 292,396,315; Median UMI counts per cell (RNA): 5,600; Median genes per cell (RNA): 2,583; Sequenced read pairs (ATAC): 246,014,072; Median fragments per cell (ATAC): 8,700.

The Cell Ranger ARC (v2.0.0) pipeline was used to process sequenced libraries with default parameters unless otherwise noted. Demultiplexed FASTQ files were generated by cellranger-arc mkfastq from BCL files. Reads were aligned to custom-built mouse (GRCm38/mm10) reference genome modified to include tdTomato using cellranger-arc count. Reads with de-duplicated and valid cell barcodes were used to build gene-by-barcode (scRNA-seq) and peak-by-barcode (scATAC-seq) matrices by cellranger-arc count per genotype. Individual matrices were aggregated to a single feature-barcode matrix file containing every genotype using cellranger-arc aggr without depth normalization (--normalize=none).

#### scRNA-seq data analysis

*Seurat:* An aggregated feature-barcode matrix was used as input to Seurat^110^ (v4.0.5, https://satijalab.org/seurat/) in R (v4.1.1, https://cran.r-project.org/). After imputing missing values to zero in metadata, outlier removal was performed on the number of counts per gene and percent reads mapping to mitochondrial genome (mitochondrial percentage). Lower limits for the number of counts per gene and mitochondrial percentage were set to 100 counts per gene and three standard deviations (SD) below the mean, respectively. Upper limits were set to three SD above the mean for both metrics. Negative datapoints created by subtraction of three SD from the mean were reset to 1, while datapoints that exceeded the upper limits were reset to the maximum datapoint. Cells were removed if they were more extreme then the upper/lower limits, or if they were eliminated from the scATAC-seq dataset during QC. The numbers of remaining cells were following: WT E12.5: 6,391; WT E15.5: 11,477; Het E12.5: 6,608; Het E15.5 10,027; KO E12.5: 8,546; KO E15.5: 8,607. Remaining cells were proceeded to the normalization workflow using Seurat::SCTransform using default parameters. For integration of scRNA-seq datasets from E12.5 and E15.5, 3,000 variable features were found using Seurat::SelectIntegrationFeatures on SCTransformed data. Prior to integration, anchors were identified using Seurat::FindIntegrationAnchors with the parameters dims, anchors.features, and normalization.method set to 1:30, the 30,000 variable features, and SCT, respectively. The integration was performed using Seurat::IntegrateData with identical dims and normalization.method to those from Seurat::FindIntegrationAnchors, along with the computed anchors. Dimensionality reduction was performed using Seurat::RunPCA and Seurat::RunUMAP with the parameters dims and umap.method set to 1:25 and uwot respectively on the integrated data.

#### scATAC-seq data analysis

*Signac:* An aggregated peak-by-barcode matrix was used as input to Signac^111^ (v1.4.0, https://stuartlab.org/signac) pipeline in R (v4.1.1). After imputing missing values in metadata as described above, outlier removal was performed on the number of chromatin accessibility peaks, transcription start site (TSS) enrichment score, and mitochondrial percentage. Lower limits were set to 1,000 counts per feature for the number of peaks and 2 for TSS enrichment score. Lower limit for mitochondrial percentage and all the upper limits were determined as described in the scRNA-seq analysis. Cells were removed if they were more extreme than the lower/upper limits in individual metrics, or if they were eliminated in corresponding scRNA-seq dataset during QC. Peaks were normalized via Term Frequency–Inverse Document Frequency (TF-IDF) method using Signac::RunTFIDF with default parameters. Highly variable peaks were found by Signac::FindTopFeatures with the min.cutoff set to 10. Dimensionality reduction was performed using Signac::RunSVD with default parameters. For integration of scATAC-seq datasets from E12.5 and E15.5, anchors were identified using Seurat::FindIntegrationAnchors with the parameters anchor.features and dims set to all features and 2:30, respectively. Integration of two scATAC-seq datasets was conducted using Seurat::IntegrateEmbeddings taking advantage of reciprocal LSI projection (RLSI), as instructed by Signac standard workflow. The Seurat::IntegrateEmbeddings ran using the previously computed anchors and 1 to 30 dimensions.

#### Multimodal analysis

*Seurat:* Multimodal data integration was started by finding Weighted Nearest Neighbor (WNN)^72^ using Seurat::FindMultiModalNeighbors on scRNA-seq and scATAC-seq datasets with or without age-specific integration along with reduction lists set to PCA (1 to 25 dimensions) for scRNA-seq and RLSI (1 to 20 dimensions) for scATAC-seq^71^. Subsequent dimensionality reduction was performed using Seurat::RunUMAP on weighted.nn with default parameters. Cells were clustered on the WNN graph with Leiden algorithm and resolution 0.8 using Seurat::FindClusters. To visualize the DEGs between populations subsetted using Seurat, we employed the EnhancedVolcano tool. For significant determination, parameters were set to pCutoff = 10e^-6^ and FCcutoff = 0.2. By default, the UMAP from the age-integrated multimodal analysis was utilized throughout the study. The expression of *Sst* and *Pde1a* genes in was examined on a multimodal UMAP at E15.5 without age integration.

For QC, the TSS enrichment score and the approximate ratio of mononucleosomal to nucleosome-free fragments (nucleosome signal) were computed using the functions Signac::TSSEnrichment and Signac::NucleosomeSignal with default parameters, respectively. The mitochondrial percentage was computed using the function Seurat::PercentageFeatureSet, which matches the pattern of gene names to “^[Mm][Tt]-“.

#### Differential analysis via DA-seq

Single cell DA analysis was conducted using DA-seq^74^ (v1.0.0, https://klugerlab.github.io/DAseq) in R (v4.1.1). The DA-seq used a PCA matrix as input, which was computed by Seurat after the multimodal integration of age-integrated scRNA-seq and scATAC-seq datasets. Cells were projected onto 2D space based on UMAP coordinates, which were computed by Seurat after multimodal integration of age-integrated scRNA-seq and scATAC-seq datasets. DA cells were determined by running DAseq::getDAcells, with the k.vector parameter set to every 50 between 50 and 500, and DAseq::updateDAcells with the pred.thres parameter set to +/-0.7. DA-seq conducted a random permutation test on abundance scores, using a threshold of +/- 0.7, to identify cells with an abundance score greater than 0.7 or less than -0.7.

#### Visualization of Multiome data

UMAP coordinates and WNN clustering, computed by Seurat on multimodal integrated datasets with or without age-specific integration, were imported to the Loupe Browser (v6.0.0, 10x Genomics). The expression of genes of interest and multimodal clustering were visualized using the Loupe Browser. DA analysis was visualized on a UMAP created using the functions DAseq::getDAcells and DAseq::updateDAcells in R, which take advantage of ggplot2 (v3.3.5). UMAPs visualizing the genotype and age in the DA analysis were generated using the Seurat::DimPlot. For QC, the following metrics were extracted from the metadata of the Seurat object: the number of ATAC fragments (nCount_ATAC), nucleosome signal (nucleosome_signal), the TSS enrichment score (TSS.enrichment), the number of RNA reads (nCount_RNA) and mitochondrial percentage (percent.mt). These metrics were then visualized using violin plots with the ggplot2 package in R.

To infer the developmental trajectories of E12.5 and E15.5 MGE cells across all conditions, Slingshot^112^ was used to generate trajectories onto UMAP projections. The Slingshot function was applied to the WNN matrix as described above, enabling the identification of neural developmental progression along the inferred trajectories.

### Statistics and reproducibility

#### Cell counts

All cell counts were performed by hand and blind to genotype. Number of brain sections per mice and mice per genotype are described above and in figure legends. One-way ANOVA was used to compare WT, Het and KO for all brain regions, followed by Tukey’s Multiple Comparison Test to identify significant differences between conditions. All statistical analysis was performed on GraphPad Prism (version 9.4.1). All raw cell counts, ANOVA F- and P-values, and results of Tukey’s multiple comparison tests related to Figures 2-4, Figure 6, Supplementary Figures 1-3 and Supplementary Figure 5 can be found in the Source Data file.

#### Electrophysiology

Collection of data and analysis were not performed blind and were non-randomized. Data from electrophysiological recordings are presented throughout as mean ± s.e.m unless otherwise noted. Unless indicated, all statistical comparisons were non-parametric. Number of recorded neurons (n) and number of animals (N) used were reported for each figure. All data were analyzed using pClamp10, GraphPad Prism and Neurolucida 360 software.

#### Sample size determination

No statistical method was used to predetermine sample size. For Western Blots, we desired n ≥ 2 biological replicates for each genotype to determine the percent of decreased H3K27me3 signal in the *Ezh2* KO. For cell counts with *Nkx2.1-Cre* mice, we wanted n ≥ 5 mice per genotype from at least 3 different litters. For cell counts with *Dlx5/6-Cre* mice, we wanted n ≥ 4 WT and KO mice per genotype from at least 3 different litters. For electrophysiological recordings, we wanted to record from ≥ 10 neurons for each condition. For the single cell Multiome experiments, we wanted a minimum of 5,000 high-quality sequenced nuclei per condition (age and genotype), which should be sufficient to identify significant differences between conditions. This goal required ∼15,000 nuclei input for each sample (with the expectation of recovering ∼30-70% of nuclei/cells for each reaction based on 10x Genomics recommendations and our previous experience). Viable nuclei that passed QC ranged from 6,391-11,477 nuclei per condition (see above). Per standard single cell sequencing protocols, nuclei that did not pass stringent QC measurements (nCount_RNA and % mitochondria reads for snRNA; nCount_ATAC, nucleosome_signal and TSS enrichment for snATAC) in the Multiome datasets were excluded from analysis (as detailed in Supplementary Figure 6). For the CUT&Tag experiments, we strove for 100,000 nuclei for each reaction (actual range from 90,000-120,000 per reaction), with n = 3 biological replicates for each condition (age and genotype). All computational and statistical analysis are discussed in detail above and/or in the legends of the relevant figures and tables. All attempts at replication were successful.

## Supporting information

Supplementary Table 1_E12Multiome

Supplementary Table 2_E15Multiome

Supplementary Table 3_E12CUTTAG

Supplementary Table 4_E15CUTTAG

## ACKNOWLEDGEMENTS

We thank S. Coon, F. Faucz, J. Iben, T. Li and other members of the NICHD Molecular Genomic Core; members of the Petros Lab for helpful discussions and feedback on the manuscript. Further information and requests for resources and reagents should be directed to and will be fulfilled by the Lead Contact, Timothy J. Petros. This work was supported by *Eunice Kennedy Shriver* NICHD Intramural Awards to T.J.P., P.P.R. and R.K.D.; NIMH Intramural Award to S.L.; an NICHD Scientific Director’s Award to T.J.P. and P.P.R.

## DATA AVAILABILITY

All sequencing data (raw and processed files) generated in this study has been deposited in the Gene Expression Omnibus (GEO) database with the following accession numbers: <u>GSE233190</u> (CUT&Tag dataset), <u>GSE233153</u> (single nuclei Multiome dataset, which includes the snRNA/GEX <u>GSE233151</u> and the snATAC <u>GSE233152</u> data). No custom code was used in the manuscript, and all computational pipelines are described in the methods. Please contact the corresponding author for more information if needed.

## AUTHOR CONTRIBUTIONS

C.T.R and T.J.P. designed the study and wrote the paper. C.T.R., D.A., Y.Z. and T.J.P. extracted and purified nuclei. C.T.R and D.A. performed CUT&Tag experiments. Y.Z. prepared single cell Multiome sequencing libraries. D.A., L.E. and Y.Z performed immunostaining, microscopy and cell counts. C.T.R., D.A, M.A., D.R.L., P.P.R and R.K.D. performed bioinformatic analysis on Multiome and/or CUT&Tag data. S.N. and S.L. performed electrophysiology experiments and analysis. P.S. performed morphological reconstructions. P.P.R., R.K.D., S.L. and T.J.P. supervised the project.

---

**Supplementary Figure 1.**
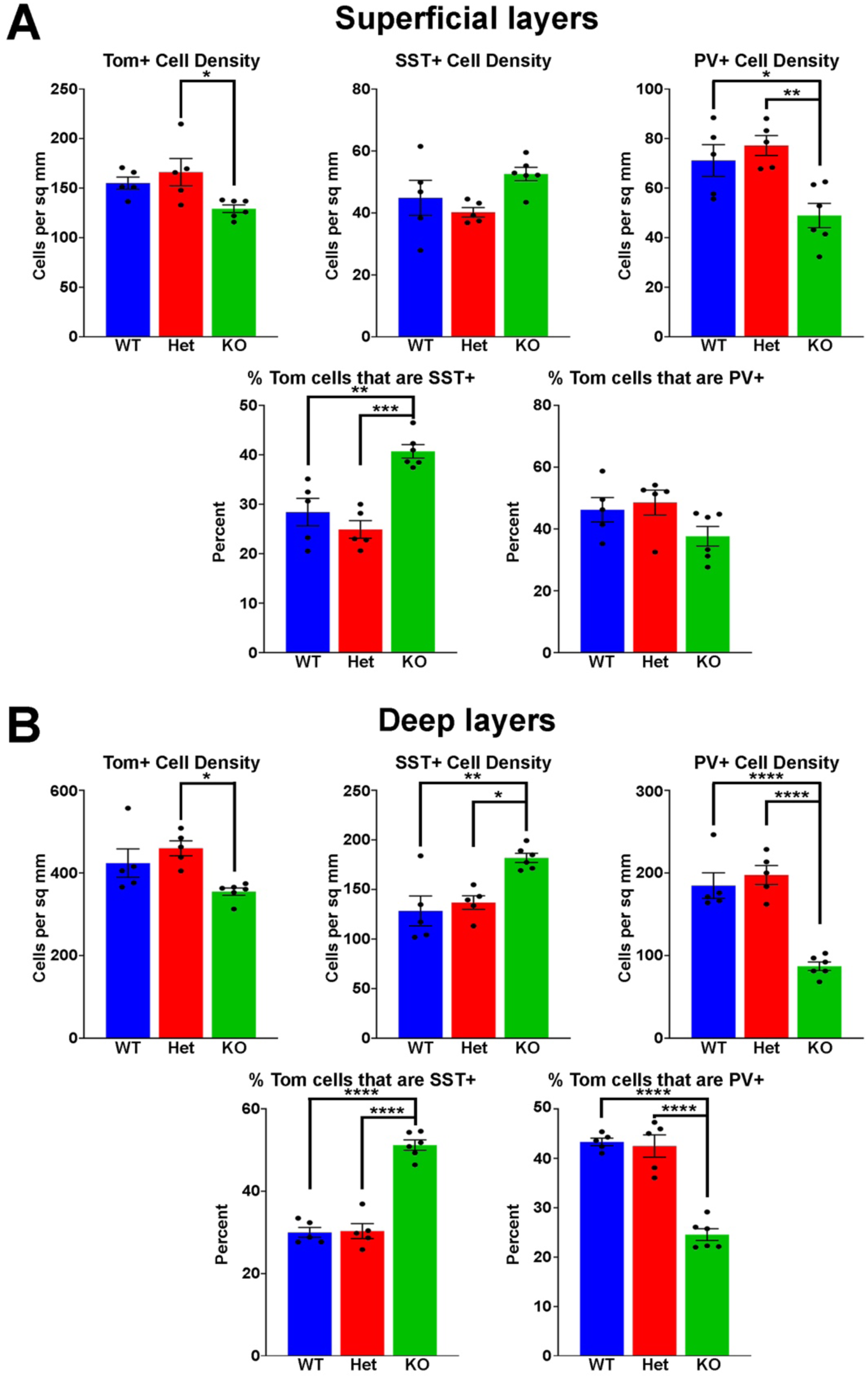
Comparison of SST+ and PV+ interneurons in superficial and deep cortical layers. **A-B.** Graphs displaying the density of Tom+, SST+ and PV+ cells (top), and percent of Tom+ cells expressing SST or PV (bottom), for superficial (**A**) and deep (**B**) cortical layers. All stats are one-way ANOVA followed by Tukey’s multiple comparison tests: * = p ≤ .05, ** = p ≤ .005, *** = p ≤ .0005, **** = p ≤ .0001. n = 5 WT, 5 Het and 6 KO brains, from 4 different litters.

**Supplementary Figure 2.**
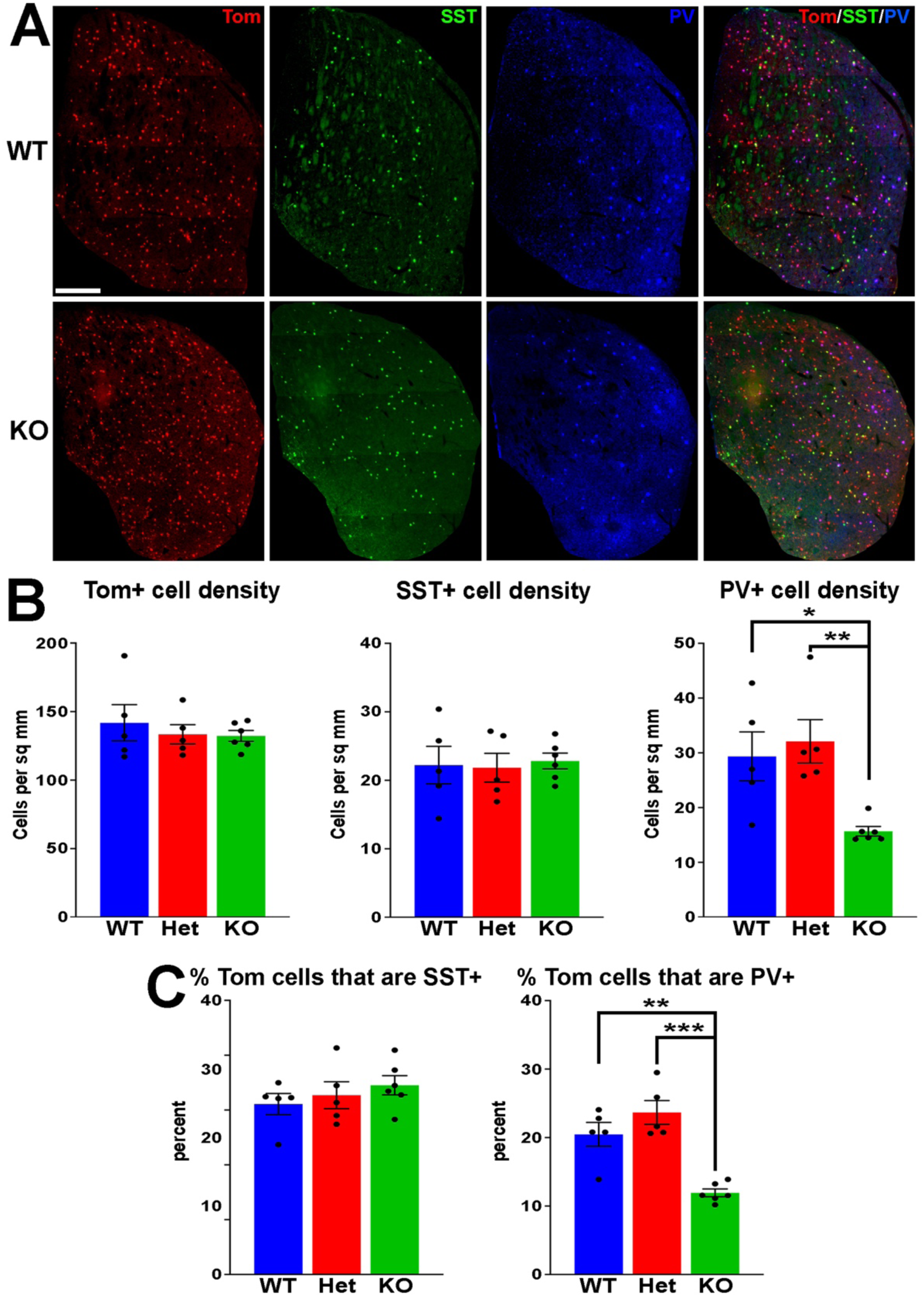
Changes in striatal interneuron fate in *Ezh2* KO mice. **A.** Representative images through the striatum of P30 *Nkx2.1-Cre*;*Ezh2*;*Ai9* WT and KO mice stained for SST (green) and PV (white/blue). Scale bar = 500 μm. **B.** Graphs displaying the density of Tom+, SST+ and PV+ cells (*top*) and the percent of Tom+ cells expressing SST or PV (*bottom*). All stats are one-way ANOVA followed by Tukey’s multiple comparison tests: * = p ≤ .05, ** = p ≤ .005, *** = p ≤ .0005. n = 5 WT, 5 Het and 6 KO brains, from 4 different litters.

**Supplementary Figure 3.**
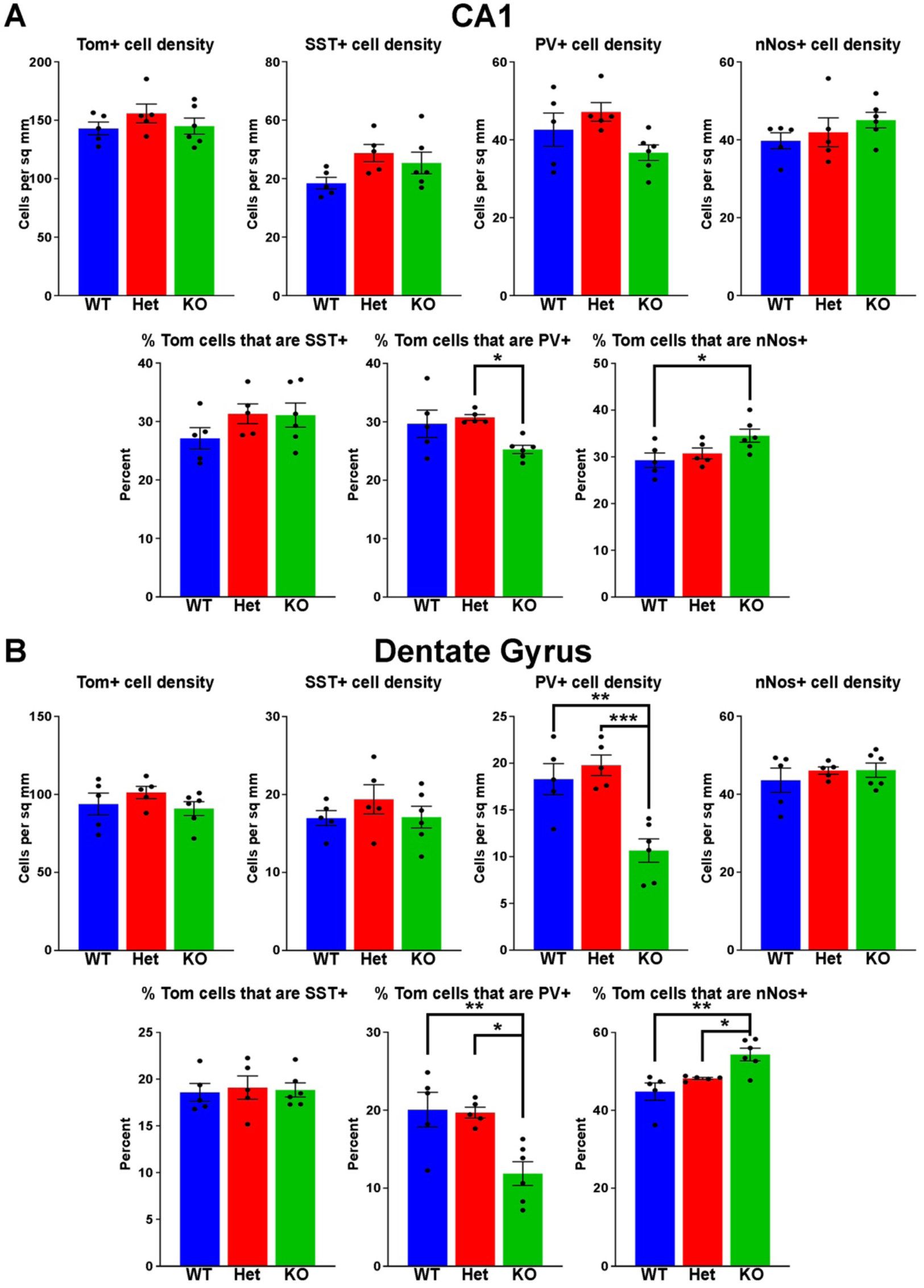
Cell fate changes in the CA1 and dentate gyrus of *Ezh2* KO mice. **A.**Graphs displaying the density of Tom+, SST+, PV+ and nNos+ cells (top) and the percent of Tom+ cells expressing SST, PV or nNos (bottom) in the CA1 region of the hippocampus. **B.** Graphs displaying the density of Tom+, SST+, PV+ and nNos+ cells (top) and the percent of Tom+ cells expressing SST, PV or nNos (bottom) in the dentate gyrus region of the hippocampus. All stats are one-way ANOVA followed by Tukey’s multiple comparison tests: * = p ≤ .05, ** = p ≤ .005, *** = p ≤ .0005. n = 5 WT, 5 Het and 6 KO brains, from 4 different litters.

**Supplementary Figure 4.**
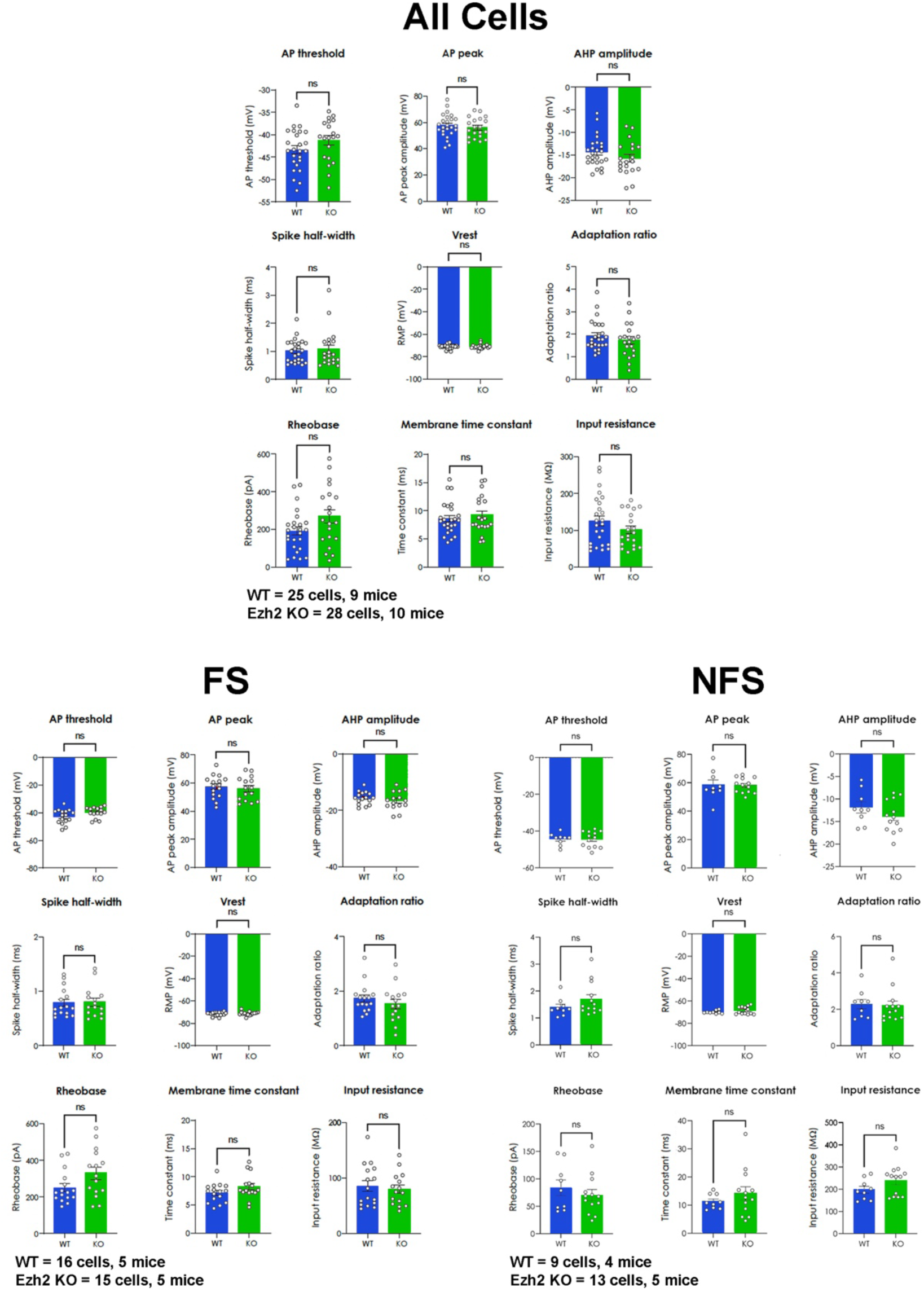
No differences in intrinsic physiology properties of MGE-derived cortical interneurons in *Ezh2* KO mice. Graphs depicting intrinsic properties of layer V/VI Tom+ cortical interneurons from WT and KO cells from fast-spiking (FS, *lower left*), non-fast-spiking (NFS, *lower right*) and combined FS and NFS cells (*top*). All statistics are Mann-Whitney test. WT = 16 FS cells from 6 mice and 9 NFS cells from 4 mice; KO = 15 FS cells from 5 mice and 13 NFS cells from 5 mice.

**Supplementary Figure 5.**
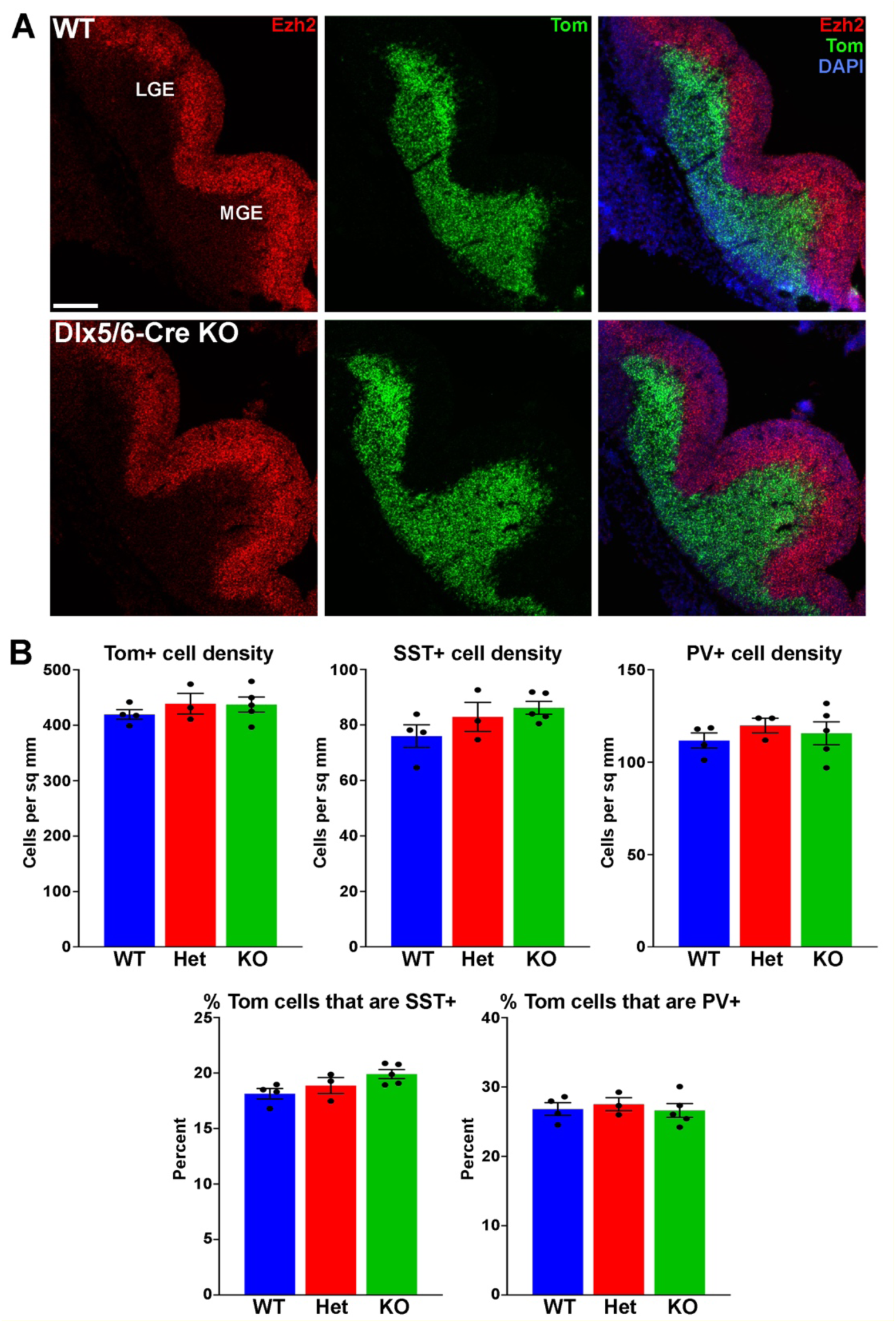
Normal interneuron densities and subtype distributions in the cortices of *Dlx5/6-Cre*;*Ezh2* KO mice. **A.** *In situ* hybridizations of *Ezh2* (red) and *tdTomato* (green) in the E12.5 MGE of *Dlx5/6-Cre*;*Ezh2*;*Ai9* WT and KO mice. Scale bar = 200 μm. **B.** Graphs displaying the density of Tom+, SST+ and PV+ cells (top) and the percent of Tom+ cells expressing SST or PV (bottom) in the cortex of WT, Het and *Dlx5/6-Cre* KO mice. All stats are one-way ANOVA followed by Tukey’s multiple comparison tests. n = 4 WT, 3 Het and 5 KO brains, from 3 different litters.

**Supplementary Figure 6.**
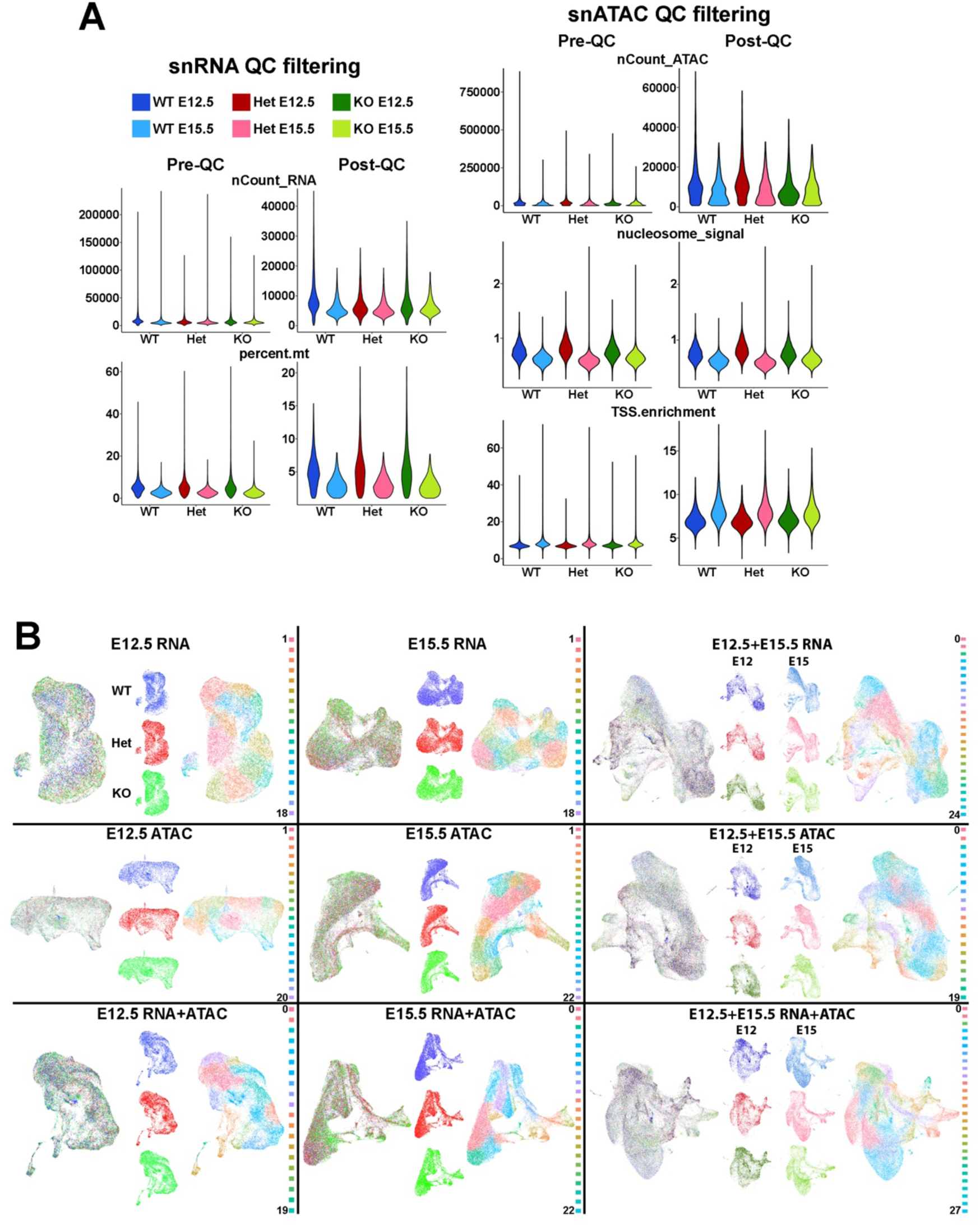
Single cell Multiome data from *Ezh2* WT and KO mice. **A.** Visualization of quality control (QC) metrics before (Pre-QC) and after (Post-QC) filtering of outliers. The number of RNA reads (nCount_RNA), mitochondrial percentage (percent.mt), the number of ATAC fragments (nCount_ATAC), nucleosome signal (nucleosome_signal), and the TSS enrichment score (TSS.enrichment) are shown. **B.** UMAP plots of Multiome data separated by age (E12.5, E15.5 and combined) and modality (RNA, ATAC and integrated RNA+ATAC), with putative cell clusters. WT = blue, Het = red, KO = green.

